# Nonlinear responses to temperature and precipitation shape the distribution of *Aedes sierrensis* in North America

**DOI:** 10.64898/2026.07.28.741332

**Authors:** Erin Mordecai

## Abstract

Understanding the ecological determinants of species ranges is a central goal of ecology. Novel tools like global datasets and machine learning models allow us to describe species ranges with increasing scope and accuracy, and to develop and test ecological hypotheses about their determinants. Here, I focus on the widespread nuisance mosquito and dog heartworm vector *Aedes sierrensis*, a tree hole-breeding mosquito that is widespread and abundance within its native range in western North America, and develop species distribution models (SDMs) to characterize the species range and its environmental determinants. I find that the species range is highly predictable from long-term average bioclimatic variables. Temperature and precipitation were the primary determinants: suitability was highest at wet-season average temperatures of 0 – 10°C, minimum temperatures of −5 – 5°C, and summer temperatures of 8 – 22°C in environments with adequate seasonal rainfall concentrated in the winter. After accounting for climate, land cover variables showed minimal importance for prediction, but suitability was higher in forests and outside of urban areas. The results are consistent with ecological knowledge of the *Ae. sierrensis* life cycle from field observations and previous laboratory experiments, suggesting that individual physiological constraints scale up to determine species distributional limits.

## Introduction

Climate and habitat requirements determine the distributions of living organisms (Elton 1927, Hutchinson 1957). All living things require suitable biophysical conditions that permit physiological performance, including temperatures that foster growth, development, survival, and reproduction, as well as physical habitat and biotic and abiotic resources such as food, water, and nutrients. Biotic interactions can further facilitate (in the case of mutualism) or limit (in the case of competition, predation, or parasitism) population growth, thereby shaping species distributions. These ecological niches can be static or dynamic, fully occupied or partially unfilled (Pulliam 2000, Matthiopoulos 2022). Taken together, the complexity of environmental conditions that shape organismal distributions and population trajectories complicate identifying the ecological factors that determine a species range, particularly as species may be actively expanding (in the case of invasive species), contracting (in the cases of threatened species), or shifting in response to environmental change and anthropogenic pressure.

The last century and a half have been defined by anthropogenic change. While humans have always modified their environment, change has accelerated since the industrial revolution, including rapid land use change from natural ecosystems to managed and converted land cover types (accelerated by mechanized equipment), anthropogenic climate change driven by increases in pollution and greenhouse gas emissions from industrial and intensified agricultural production, and increased global trade and travel, which facilitates biotic exchange of non-native species and human commensals that alter species interactions (Foley et al. 2005, Rockström et al. 2009). In an era of rapidly changing climate and land use, it is critical to understand how global change is affecting the distributions and population trajectories of species. This first requires understanding the fundamental ecological requirements of each species. This is particularly challenging as environmental features like climatic regimes and biotic regimes (e.g., ecoregions) are often tightly correlated, making it difficult to parse out the role of distinct climatic and other ecological features.

Species distribution models (SDMs) are often used in ecology to delineate the geographic boundaries and ecological requirements of species by extrapolating from their currently observed occurrence records (Elith et al. 2006, 2008, Barker and MacIsaac 2022, Lippi et al. 2023, Singleton et al. 2024). SDMs are a family of modeling approaches that can use a variety of algorithms and environmental features, but the key concept is to use flexible functions to identify combinations of variables that differentiate occurrences from absences or ‘background’ points, distinguishing where a species is present from where it could have been sampled but was not. These models can be useful for formally predicting species distributional boundaries and for identifying ecological relationships that may delimit species ranges. However, as an observational modeling approach, SDMs can also be sensitive to model assumptions, algorithm choice, occurrence and background sampling choices, variable inclusion, and more (Barker and MacIsaac 2022, Lippi et al. 2023, Singleton et al. 2024). Thus, it is often helpful to fit and compare several models rather than relying on a single model specification, and field- or lab-based validation of species ecological requirements is important for understanding mechanisms.

Although SDMs fundamentally assume that where a species currently occurs reflects its ecological requirements—in other words, that species are at equilibrium with respect to their environmental constraints—SDMs are often used in contexts of change: invasive species, threatened or endangered species, or species that are noxious to humans and subject to control measures (Araújo et al. 2019, Barker and MacIsaac 2022, Lippi et al. 2023). Here, I develop SDMs for a species that is geographically widespread, abundant, confined to its native range, and not actively undergoing decline: the western tree hole mosquito, *Aedes sierrensis* (Ludlow 1905), which is distributed throughout western North America (Darsie and Ward 2005). The goal of the SDMs is to understand the species current distribution and the environmental variables that best predict its limits. Unlike many other mosquito species for which SDMs have been developed (Barker and MacIsaac 2022, Lippi et al. 2023), *Ae. sierrensis* is not invasive and is not a vector of human disease. As a nuisance mosquito and a vector of the parasite *Dirofilaria immitis*, which causes dog heartworm (Ledesma and Harrington 2011, Couper and Mordecai 2022), it is a target of vector surveillance and control mainly where it overlaps with human residential areas. It has also been developed as a model system for studying climate adaptation and host – parasite interactions, making it scientifically and ecologically relevant (Washburn et al. 1988, 1989, Ismail et al. 2023, Couper et al. 2024a, Lyberger et al. 2024, Couper et al. 2025, Farner et al. 2026). Understanding the ecological determinants of its distribution is useful for basic ecological knowledge, and downstream for applied vector control and global change biology. As a non-invasive, non-threatened species that is not subject to major anthropogenic modifications such as habitat loss or aggressive and widespread vector control, I expect that the species range will be highly predictable—because it is close to its environmental equilibrium—and constrained by environmental variables such as climate and land cover.

The *Ae. sierrensis* life cycle is adapted to the climate of western North America and the phenology of water-filled tree holes (phytotelemata) in which it develops. Unusually for mosquitoes, it is univoltine, producing a single generation per year. It lays its eggs in tree holes that become filled with water during the wet winter season; larvae undergo development in the tree holes and persist for as long into the spring as the tree holes hold water, typically spending 2-6 months and entering diapause at the final (L4) larval stage, then pupating and emerging as adults when tree holes begin to dry down in the spring and early summer. Emerging adults mate and females blood feed on mammalian hosts and lay eggs in tree holes, typically near their natal tree hole because they are weak fliers. Eggs remain dormant and desiccation resistant in diapause through the summer and fall until rain fills the tree holes in late fall or early winter.

In this work, I develop SDMs for *Ae. sierrensis*, to ask: (1) what is the species’ most likely geographic range, and how predictable is it? (2) which climate and land cover variables are most predictive of this range? (3) does the species range respond to environmental gradients with smooth, gradual responses or sharp thresholds? (4) can different combinations of environmental variables produce similar model predictions, due to their underlying correlations? and (5) are predicted environmental responses consistent with laboratory-measured thermal performance? I develop a set of candidate models based on evaluating the ecological hypotheses that the *Ae. sierrensis* distribution is limited to highly seasonal environments with cool wet seasons and hot dry seasons (e.g., Mediterranean climates) and vegetated areas with trees that can serve as immature habitat, and further limited by extreme summer highs and winter lows and sufficient rainfall to sustain several months per year in which water-filled tree hole immature habitat is available.

As the results show, the species distribution is highly predictable using a suite of overlapping features, including temperature and precipitation means and seasonality, while land cover contributes minimally to prediction accuracy, likely because its variation is already absorbed by climate variables or because coarse land cover classes do not capture tree hole habitat availability.

## Methods

### Overview

The goal of this model is to predict and explain the current distribution of *Ae. sierrensis* across its full geographic extent in North America (USA, Canada, and Mexico). A further aim is to evaluate which environmental variables hold the most predictive power, and whether these correspond to aspects of the mosquito’s biology, suggesting potential causal relationships. I further use the model to test hypotheses about the climatic and land cover determinants of *Ae. sierrensis* range limits (as outlined in the Introduction). To meet these objectives, I fit a series of candidate models that capture different aspects of the environmental space while limiting correlations among variables included.

Throughout the modeling, I checked that reproducibility standards and best practices (Araújo et al. 2019, Feng et al. 2019, Zurell et al. 2020, Barker and MacIsaac 2022) were followed as closely as possible. This includes reporting through the ODMAP (Overview, Data, Model, Assessment, and Prediction) framework (Appendix S3).

### Study system

The western tree hole mosquito, *Aedes sierrensis* (Ludlow 1905), is broadly distributed across western North America in coastal and inland areas that have seasonal precipitation concentrated in the winter and spring. Although *Ae. sierrensis* is widespread and common across western North America, it has a distinct geographic distribution limited to within a few hundred kilometers of the Pacific coast (Darsie and Ward 2005). This suggests that it has distinct climatic requirements associated with coastal climate influence.

Adult females primarily lay their eggs in naturally occurring tree holes or tree stumps common in oak species such as live oaks (*Quercus agrifolia* and *Q. wislizeni*), black oaks (*Q. kelloggii*), blue oaks (*Q.douglasii*), and valley oaks (*Q. lobata*), as well as some conifers. *Ae. sierrensis* larvae hatch when tree holes fill with water during fall, winter, or spring rains or during snow melt in cooler areas, and progress through four larval stages (L1-L4) before pupating then emerging as adults as tree holes dry down in the late spring and summer. Adults mate and females seek blood meals from mammalian hosts before laying their eggs in tree holes, completing the life cycle. Because the univoltine (one generation per year) *Ae. sierrensis* life cycle can include two diapauses—at the egg and L4 larval stages—the mosquito spends most of the year in immature stages and only a few weeks as an adult. At larval stages, *Ae. sierrensis* graze on bacteria, algae, and microorganisms in the tree hole water, including *Lambornella clarki*, a ciliate that facultatively parasitizes and kills *Ae. sierrensis* larvae—the primary biotic interaction of importance in the mosquito’s life cycle (Washburn et al. 1991).

Despite decades of research on the species, this work develops the first published SDMs for *Ae. sierrensis*. Previous experimental work at constant temperatures has demonstrated that thermal performance and limits vary among traits in the *Ae. sierrensis* life cycle, ranging from lower thermal limits of 0.18 – 7.16°C to thermal optima of 14.25 – 27.22°C to upper thermal limits of 31.60 – 34.00°C (Couper et al. 2024a). These thermal responses were consistent across populations spanning 1,200 km (10° latitude) except for larval and pupal development rates, for which the upper thermal limits varied by 1.6°C across this geographic range (Couper et al. 2024a). A previous study (Couper and Mordecai 2022) modeled *Ae. sierrensis* abundance from vector surveillance data and weather conditions matched to the sampling period, but did not report the inferred geographic range, environmental predictors, and environment – suitability relationships because the *Ae. sierrensis* sub-model was one of many mosquito models developed as inputs into a model of dog heartworm incidence. Robust field surveillance programs by California, Oregon, Washington, and Utah vector control agencies have documented peak abundance of adult *Ae. sierrensis* in April - June, though adult mosquitoes have been detected as late as October (Couper et al. 2024a)(VectorSurv).

### Candidate models

Here, the goal was to produce an accurate suitability map with interpretable environmental response relationships, testing hypotheses about important temperature, rainfall, and land cover relationships, rather than to obtain the highest possible predictive accuracy by maximizing variable inclusion. Candidate models (Table 1) encapsulating these hypotheses include: model 1 (‘baseline’), which includes the annual averages and seasonal variation (i.e., seasonality) of temperature and precipitation; model 2 (‘seasonal averages’), which includes seasonal averages of temperature and precipitation; model 3 (‘extremes’), which includes temperature and precipitation variability and extremes; model 4 (‘hypotheses’), which most closely reflects my ecological hypotheses and includes temperature and precipitation averages during specific biologically relevant windows; and model 5 (‘kitchen sink’), a maximalist model that includes a wider range of plausibly relevant bioclimatic variables. Each of models 1-5 also includes six relevant land cover variables (forest, shrub, grassland, wetland, crop, and urban). In addition, model 6 (‘seasonal averages reduced LC’) includes the seasonal average bioclimatic variables from model 2 with only forest and urban cover (the two most relevant land cover variables); model 7 (‘hypotheses reduced LC’) includes the hypothesized biologically relevant bioclimatic variables from model 4 with only forest and urban cover; and model 8 (‘hypotheses no LC’) includes the bioclimatic variables of model 4 with no land cover variables.

**Table 1.**
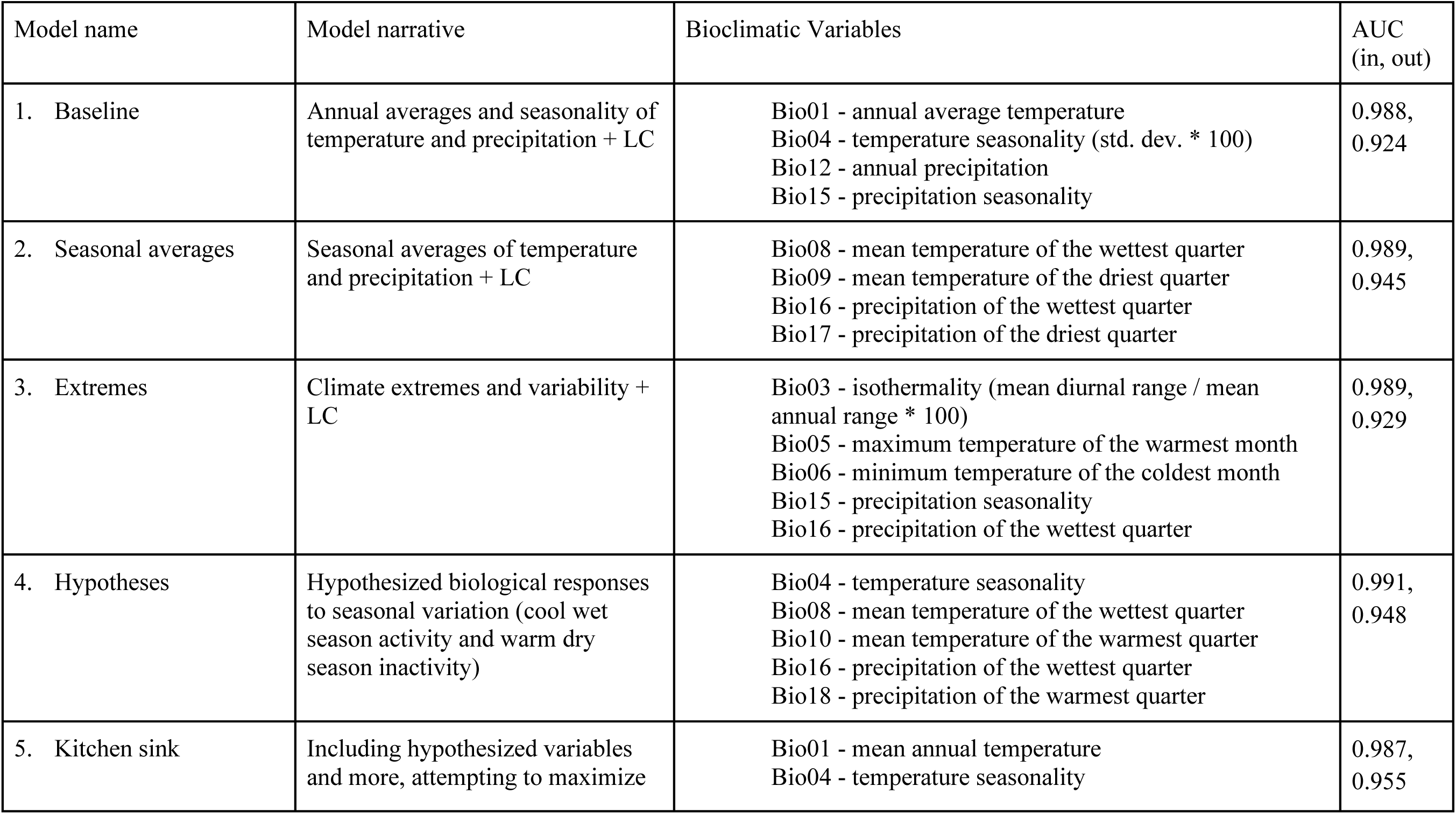
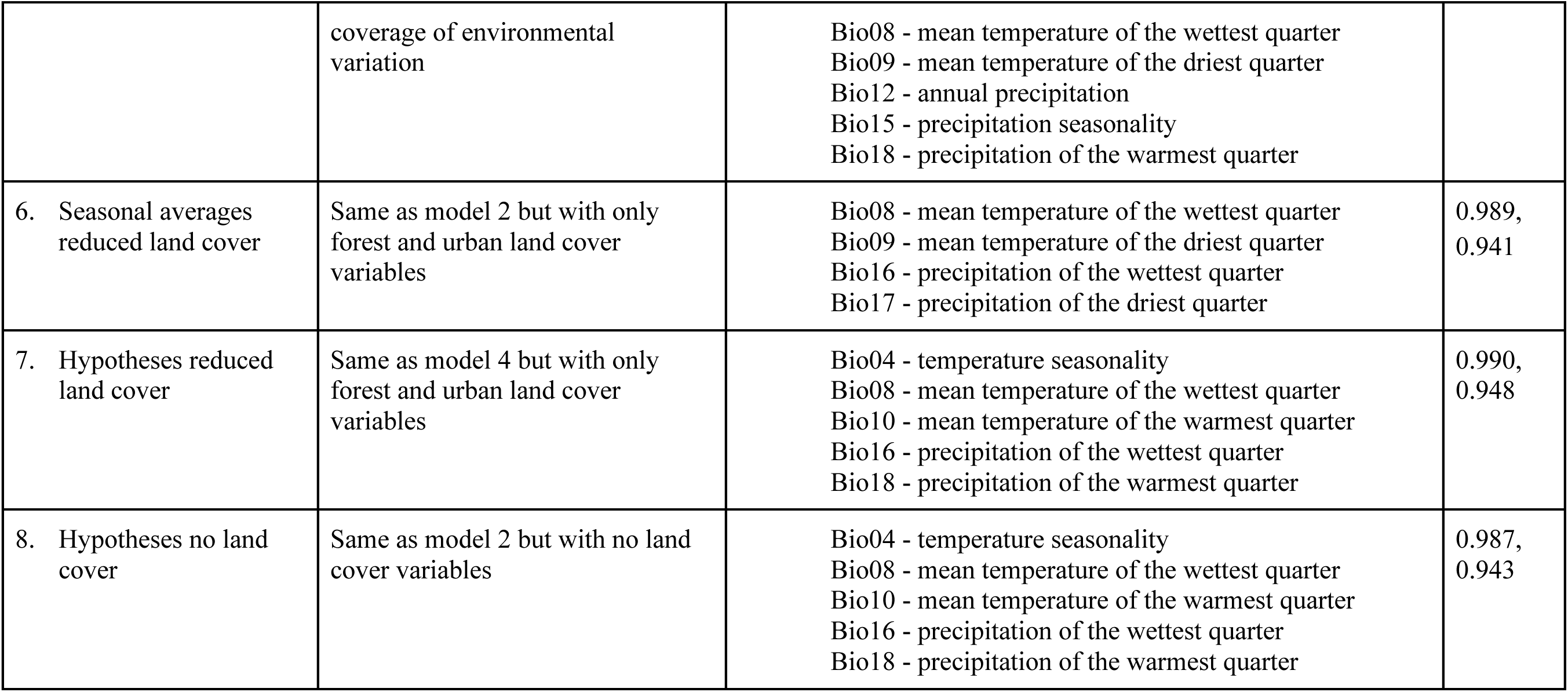
Model names, narratives, bioclimatic variables, AUC in and out of sample, and LogLoss. + LC indicates the full suite of land cover variables included: forest, urban, cropland, wetland, shrubland, grassland. Reduced LC indicates that only forest and urban land cover variables were included. No LC indicates that no land cover variables were included.

| Model name | Model narrative | Bioclimatic Variables | AUC<br>(in, out) |
| --- | --- | --- | --- |
| 1. Baseline | Annual averages and seasonality of temperature and precipitation + LC | Bio01 - annual average temperature<br>Bio04 - temperature seasonality (std. dev. * 100)<br>Bio12 - annual precipitation<br>Bio15 - precipitation seasonality | 0.988,<br>0.924 |
| 2. Seasonal averages | Seasonal averages of temperature and precipitation + LC | Bio08 - mean temperature of the wettest quarter<br>Bio09 - mean temperature of the driest quarter<br>Bio16 - precipitation of the wettest quarter<br>Bio17 - precipitation of the driest quarter | 0.989,<br>0.945 |
| 3. Extremes | Climate extremes and variability + LC | Bio03 - isothermality (mean diurnal range / mean annual range * 100)<br>Bio05 - maximum temperature of the warmest month<br>Bio06 - minimum temperature of the coldest month<br>Bio15 - precipitation seasonality<br>Bio16 - precipitation of the wettest quarter | 0.989,<br>0.929 |
| 4. Hypotheses | Hypothesized biological responses to seasonal variation (cool wet season activity and warm dry season inactivity) | Bio04 - temperature seasonality<br>Bio08 - mean temperature of the wettest quarter<br>Bio10 - mean temperature of the warmest quarter<br>Bio16 - precipitation of the wettest quarter<br>Bio18 - precipitation of the warmest quarter | 0.991,<br>0.948 |
| 5. Kitchen sink | Including hypothesized variables and more, attempting to maximize | Bio01 - mean annual temperature<br>Bio04 - temperature seasonality | 0.987,<br>0.955 |
|  | coverage of environmental variation | Bio08 - mean temperature of the wettest quarter<br>Bio09 - mean temperature of the driest quarter<br>Bio12 - annual precipitation<br>Bio15 - precipitation seasonality<br>Bio18 - precipitation of the warmest quarter |  |
| 6. Seasonal averages reduced land cover | Same as model 2 but with only forest and urban land cover variables | Bio08 - mean temperature of the wettest quarter<br>Bio09 - mean temperature of the driest quarter<br>Bio16 - precipitation of the wettest quarter<br>Bio17 - precipitation of the driest quarter | 0.989,<br>0.941 |
| 7. Hypotheses reduced land cover | Same as model 4 but with only forest and urban land cover variables | Bio04 - temperature seasonality<br>Bio08 - mean temperature of the wettest quarter<br>Bio10 - mean temperature of the warmest quarter<br>Bio16 - precipitation of the wettest quarter<br>Bio18 - precipitation of the warmest quarter | 0.990,<br>0.948 |
| 8. Hypotheses no land cover | Same as model 2 but with no land cover variables | Bio04 - temperature seasonality<br>Bio08 - mean temperature of the wettest quarter<br>Bio10 - mean temperature of the warmest quarter<br>Bio16 - precipitation of the wettest quarter<br>Bio18 - precipitation of the warmest quarter | 0.987,<br>0.943 |

I trained and compared the performance of these models through (i) spatial cross-validation (which holds out spatially structured ‘folds’ of data during model training then tests predictive performance on the held-out fold), (ii) visualizing relationships between environmental predictors and predicted environmental suitability (using partial dependence plots), and (iii) examining prediction maps compared to occurrence and background points. I then created an unweighted model average (ensemble) and visualized both the ensemble predicted suitability and the geographic locations where predictions differ among models (standard deviation of predicted suitability across models).

### Software

All analyses were conducted in R version 4.5.2 (2025-10-31) with RStudio version 2026.01.1+403 using the packages: blockcv (Valavi et al. 2025), caret (Kuhn 2008), corrplot (Taiyun 2026), data.table (Barrett et al. 2026), dismo (Hijmans et al. 2024), dplyr (Wickham et al. 2026b), forcats (Wickham et al. 2025a), future.apply (Bengtsson and R Core Team 2026), ggnewscale (Campitelli 2025), ggplot2 (Wickham et al. 2026a), mltools (Gorman 2018), mgcv (Wood 2025), patchwork (Pedersen 2025), pdp (Greenwell 2026b), PerformanceAnalytics (Peterson et al. 2026), pROC (Robin et al. 2025), purrr (Wickham et al. 2026c), raster (Hijmans et al. 2025), rBayesianOptimization (Yan 2025), RColorBrewer (Neuwirth 2022), rgbif (Chamberlain et al. 2026), rnaturalearth (Massicotte et al. 2026), rsample (Frick et al. 2026), sf (Pebesma et al. 2026), SHAPforxgboost (Liu et al. 2025), spatialsample (Mahoney et al. 2025), terra (Hijmans et al. 2026), tidyr (Wickham et al. 2025b), tidytext (Queiroz et al. 2025), tidyverse (Wickham and RStudio 2023), vip (Greenwell 2026a), and xgboost (Chen et al. 2026). Environmental data extraction was performed in Google Earth Engine (Gorelick et al. 2017) using the Google Colab python interface (Bisong 2019).

### Data

#### Occurrence records and filtering procedure

Species distribution models can estimate the probability that a species could occur in a given location as a function of environmental variables by comparing records of where the species has been detected to true absence observations or, more commonly, ‘background’ points where the species could have been sampled but was not. Accurately capturing a species ecological niche requires that the background points appropriately represent the sampling process, including potential sampling bias, that might make detection of the focal species more likely in some places than others, given that it is present. For example, sampling may be more likely to occur where there is greater human population density or research effort, and background points should capture this variation. To improve the ecological interpretability of model results, it is also important that the background points span the feasible geographic range but not too far outside of it (e.g., omitting continents and biomes where the species has never been recorded), otherwise the model can artificially inflate statistical performance simply by discriminating among these major biomes, rather than more locally in environmental space. Finally, background points should cover the feasible range of environments included within the geographic range of prediction maps. One method for accounting for these sampling biases and geographic limits is to use occurrence records of similar species as background points, aiming to capture shared bias in sampling effort (for example, if mosquitoes are less frequently sampled in remote areas, this bias in detection probability would be shared between *Ae. sierrensis* and other mosquito species). Finally, because sampling is heterogeneous in space, with abundant records in some locations and sparse records in others, it is important to filter the occurrence and background data to avoid overfitting the model on a few high-effort locations.

To meet these requirements, I downloaded occurrence records for the focal species *Ae. sierrensis* (GBIF listing: *Aedes* (*Jarnellius*) *sierrensis* (Ludlow, 1905)) and additional mosquito species that occur across North America (for use as background points): *Ae. communis*, *Ae. excrucians*, *Ae. fitchii*, *Ae. sticticus*, *Ae. triseriatus*, *Ae. vexans*, *Culex pipiens*, *Cx. tarsalis*, and *Cx. quinquefasciatus*, which together span the range of interest in the USA, Canada, and Mexico, from the Global Biodiversity Information Facility (GBIF, gbif.org) using the function ‘occ_search’ in the rgbif package. The data were downloaded on June 23, 2026. Records were included for species presences recorded between 1981 and 2025, which included latitude and longitude, and for which the basis of record was ‘HUMAN_OBSERVATION’, ‘PRESERVED_SPECIMEN’, or ‘MATERIAL_SAMPLE’; records for which the basis of record was ‘FOSSIL_SPECIMEN’ (0), ‘MATERIAL_CITATION’ (12) or ‘MACHINE_OBSERVATION’ (45) were removed. The number of records downloaded and saved after filtering for each species, along with the rationale, is summarized in Table S1. To include background points that would help to delineate the northern extent of *Ae. sierrensis*’s range, I specifically targeted mosquito species of Canada for inclusion as background points (Villeneuve et al. 2025), and likewise for the Intermountain West, USA, I searched for *Ae. vexans* occurrence points in Washington, Utah, and Idaho.

For each species, except as described in Table S1, I downloaded the first 10,000 records occurring since 1981 in North America, then dropped any records missing latitude, longitude, or year and/or falling outside of the USA, Canada, and Mexico, or falling in Hawaii. The temporal range of the points was chosen to help ensure accuracy of sampling point GPS locations and broad temporal alignment of points with environmental data sources. I visually checked all points for accuracy (using terra), noting that no points fell outside the geographic ranges of other points, in implausible environments, or in the middle of oceans. I then filtered the points using a probability mask at 1000 m resolution, selecting up to one *Ae. sierrensis* point per grid cell, then probabilistically sampled the background points (all other species) with a sampling ratio of 2 background points per 1 occurrence point. This resulted in 273 occurrence points and 546 background points (Fig. 1). This ratio balances the goals of adequately covering the range of environments in the geographic area of interest while avoiding falsely inflating model performance by overfitting on pseudo-absences.

**Figure 1.**
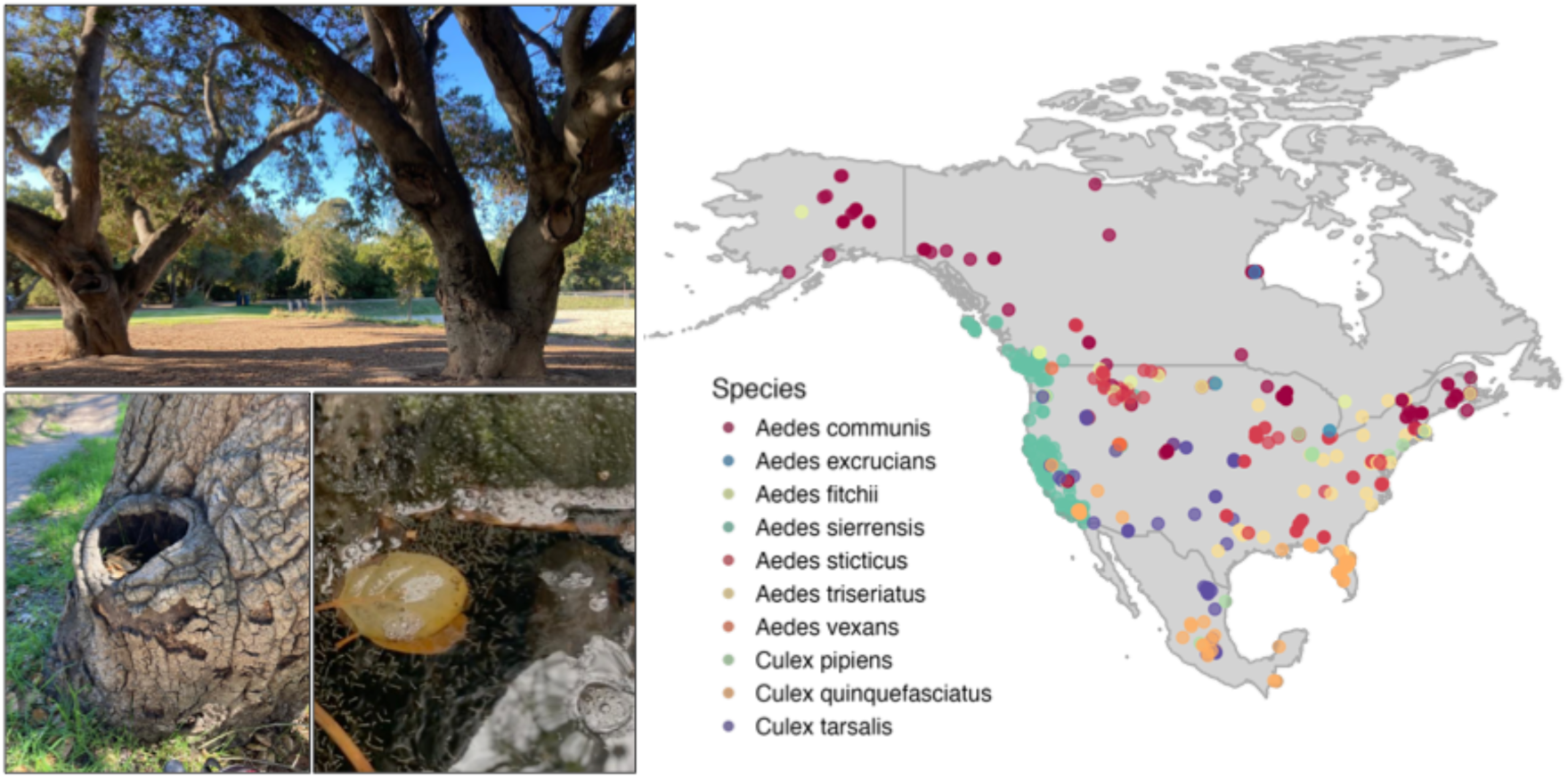
Aedes sierrensis habitat photos (left) and occurrence records, with background points, across North America (right). Photos show examples of Aedes sierrensis tree hole habitat (top: oak trees with tree holes; bottom left: a tree hole; bottom right: zoomed in view of a water-filled tree hole containing larvae; photos courtesy of Kelsey Lyberger). Map of thinned occurrence records for *Aedes sierrensis* and background points (records of other mosquito species) used to fit the models.

#### Environmental covariate data

I extracted environmental covariate data from 1 km-diameter circular buffers around each occurrence or background point, with a maximum error of 100m, in Google Earth Engine (Gorelick et al. 2017). All environmental data were downloaded on June 24, 2026.

For each buffered point, I then obtained climatology data from Worldclim version 1 (Hijmans et al. 2005) and extracted the mean of each band at 1 km resolution (bioclimatic variable names and descriptions listed in Table 1). While other databases may capture the actual weather at a given sampling point, here I am interested in how the long-term climatic conditions (as represented by bioclimatic variables) shape the probability of occurrence for *Ae. sierrensis* (Araújo et al. 2019 note that analyses using environmental data from historical time periods may better capture lagged responses of spatial distributions).

I extracted land cover data for each buffered point from the National Land Cover Database Land Cover of North America (Commission for Environmental Cooperation (CEC) et al. 2024) at 30 m resolution, selecting the modal land cover type for each buffered point. This dataset uses harmonized Landsat 8 Collection 2 Level 1 imagery from Canada from 2020, the coterminous USA 2019, Alaska from 2021, and Mexico land cover change from 2015-2020. From the 19 original land cover categories in the dataset, I reclassified them into binary categories that are ecologically relevant for mosquitoes: forest (classes 1-6), shrubland (classes 7-8), grassland (classes 9-10), wetland (class 14), cropland (class 15), urban (class 17), and water (class 18), dropping other classes that were rare or absent from the dataset. The 1 km resolution of the environmental data is consistent with the measured flight range of *Ae. sierrensis* (Yee and Anderson 1995), indicating ecological relevance. The final environmental variable list is presented in Table 1 and displayed in Figure S1.

Several bioclimatic variables had high correlations with each other (Figs. S2-S3). Although tree-based machine learning methods can handle correlated variables, including them can reduce ecological interpretability, which was the primary goal of these models. Therefore, for each model specification I selected a subset of bioclimatic variables (along with land cover variables) that represented distinct environmental variation, then compared performance among models including different variable subsets (Table 1).

### Model fitting

#### XGBoost modeling methods

To predict environmental suitability for *Ae. sierrensis*, measured as the probability of a given sampling point being the focal species versus background, and to evaluate its environmental predictors, I used extreme gradient boosted regression (XGBoost) using the package xgboost. EXGBoost is useful for this type of ecological modeling because it can capture nonlinear and interactive relationships, handle missing data, and avoids overfitting through regularization. It works by fitting a sequence of weak decision trees, each of which trains on the errors of the previous tree to iteratively refine the predictions of the model ensemble. The algorithm also uses settings to reduce overfitting, such as early stopping, shallow trees, random predictor and observation selection, and penalties for complexity, which are tuned in the fitting process.

To train and test the model, I divided the data into three spatial folds using the ‘random’ method with size = 100,000 m and 5000 iterations in the ‘cv_spatial’ function in the blockCV package (Fig. S4). Folds were stratified by presence (∼2:1 ratio of background to presence) to ensure balance among folds. I visually examined fold assignments across geographic and environmental space to further ensure balance and to check for outliers in environmental variables. All modeling procedures described below were performed iteratively, holding one fold out and training on the other two.

To tune the XGBoost model, I performed Bayesian hyperparameter optimization (BO) using the following settings: nrounds = 200, booster = “gbtree”, evaluation metric = “logloss”, objective = “binary:logistic”, nfold = 5, early_stopping_rounds = 10, init_points = 8, n_iter = 25, and searching over the following bounds: eta (0.01, 0.3), max.depth (2L, 10L), min.child.weight (1L, 15L), subsample (0.6, 1), colsample_bytree (0.6, 1).

### Assessment

#### Fit statistics, variable importance, and partial dependence plots

Logloss and area under the receiver operating curve (AUC) in- and out-of-sample were used to evaluate model performance for each held-out fold, then averaged across folds for final statistics (Table 1). All model performance statistics and predictions were consistent across the held-out folds. To evaluate feature importance—the degree to which each variable contributes to overall model predictions—I used Shapley additive explanations (SHAP) scores in the R package SHAPforxgboost. Shapley scores can be aggregated across all prediction points to evaluate an overall contribution for each variable, and they can be used in partial dependence plots, illustrating the marginal contribution of each variable to occurrence probability across all observed values of the other variables. Because Shapley scores and PDPs were similar across all folds, I aggregated across folds to calculate overall versions. I visually examined partial dependence plots to evaluate variable relationships.

### Predictions

Using the final model, I mapped the predicted occurrence probability by downloading rasters for each bioclimatic and land cover variable across the spatial extent of the USA (excluding Hawaii), Canada, and Mexico, aggregating to 1 km (for the land cover data, which had a native resolution of 30 m), and joining using a consistent coordinate reference system (CRS). Raster processing was done using the terra package in R. I input the resulting raster stack into each final XGBoost model to create a prediction surface raster. From the individual model rasters, I created an ensemble model as a simple average across the 8 candidate models (without differentially weighting models because I deemed all of them plausible), along with a map visualizing uncertainty as the standard deviation across model predictions within the ensemble.

## Results

### Model-predicted habitat suitability

*Ae. sierrensis* habitat suitability was highly predictable using publicly available occurrence and background sampling points (Fig. 1), remotely sensed environmental variables (Fig. S1, and XGBoost models. Across several model specifications that included different combinations of environmental variables, model predictive accuracy was high both in- and out-of-sample and across different held-out spatial folds (Table 1, Fig. S5). Area under the receiver operating characteristic curve (AUC, which ranges from 0 to 1 with 0.5 equivalent to predicting at random), ranged from 0.984 - 0.997 in-sample and from 0.867 - 0.980 out-of-sample across all model specifications and folds, indicating high predictive skill at discriminating occurrence from background points (i.e., habitat suitability; Fig. S5). While in-sample AUC was consistently above 0.95 across model specifications and folds, out-of-sample accuracy dipped below 0.9 only for a single fold in the baseline and extremes models, (models 1 and 3, respectively).

All models predicted high suitability in the core of the species range, along a ∼400 km wide band from the Pacific coast to inland areas from southern California, USA through coastal British Columbia, Canada, extending slightly into Baja California, Mexico, and low suitability throughout most of the rest of USA, Canada, and Mexico (Figs. 2, S6). Models predicted high to intermediate suitability in parts of the Intermountain West, USA. Several models also predicted high suitability in other parts of the geographic extent, often where there were few or no occurrence and background points, including parts of southwestern USA into highland western Mexico (models 1 and 3) and the southeastern USA (models 2, 6, and 8 and to a lesser extent models 1, 4, 5, and 7; Fig. S6). As a result, the ensemble model, a simple average of models 1-8, shows high predicted suitability in the core range along the western coast of North America from southern California to British Columbia and low suitability elsewhere (Fig. 2a). The areas of greatest uncertainty in suitability, based on the standard deviation in predicted suitability across model specifications, include the Intermountain West USA, highland western Mexico, a small pocket of the southeastern USA, and a narrow band of western Canada inland of the Pacific coast (Fig. 2b). These geographic areas of model disagreement despite high overall performance statistics highlight the importance of including a broad range of occurrence and background sampling points that fully span the environmental space included in the geographic region of interest (see Table S2 for further discussion).

**Figure 2.**
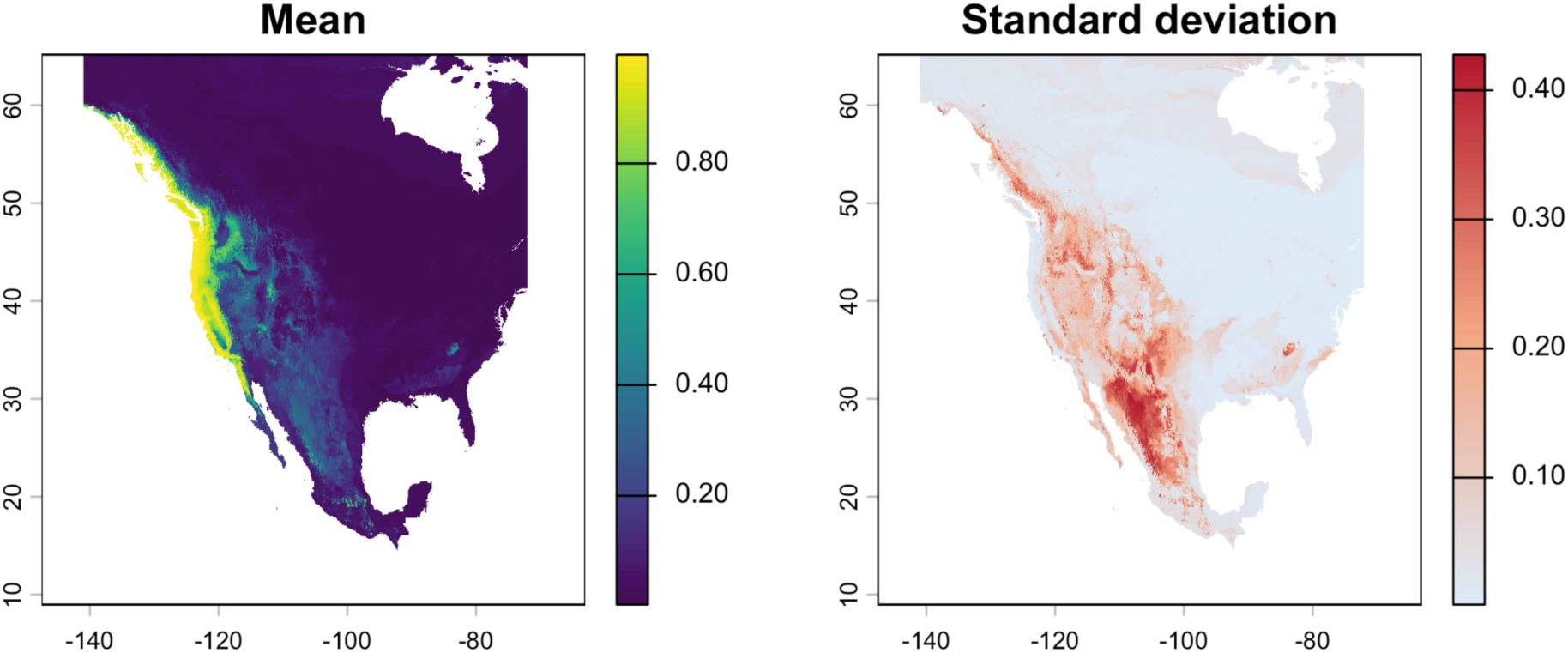
Ensemble model mean suitability prediction (left), ranging from zero (dark purple) to one (yellow), and standard deviation of predicted suitability across models included in the ensemble (right), ranging from zero (gray) to 0.4 (red). Model agreement was high in the core part of the range along the coast of western North America and in the highly unsuitable ranges of interior Canada and midwestern to eastern United States, while model disagreement was highest in the forested regions of central and western Mexico and the Intermountain West of the United States and Canada, as well as the southeastern United States.

#### Environmental drivers of suitability

Because many bioclimatic and land cover variables are correlated (Figs. S2-S3), I fit a series of models with varying combinations of environmental variables to test the hypothesis that habitat suitability for *Ae. sierrensis* is highest with high precipitation seasonality and overall rainfall, low temperature seasonality (i.e., moderate temperatures year-round), with most rain falling in the cold season, and in tree-covered areas. Multiple combinations of bioclimatic variables can potentially capture these conditions. As a result, models combining different environmental variables produced broadly similar predictive accuracy (Fig. S5) and suitability maps (Fig. S6) but differed in variable importance across models (Fig. 3). Climate variables were always more important than land cover variables, based on both SHAP scores (Fig. 3; hereafter used as the primary importance metric) and gain (Fig. S7).

**Figure 3.**
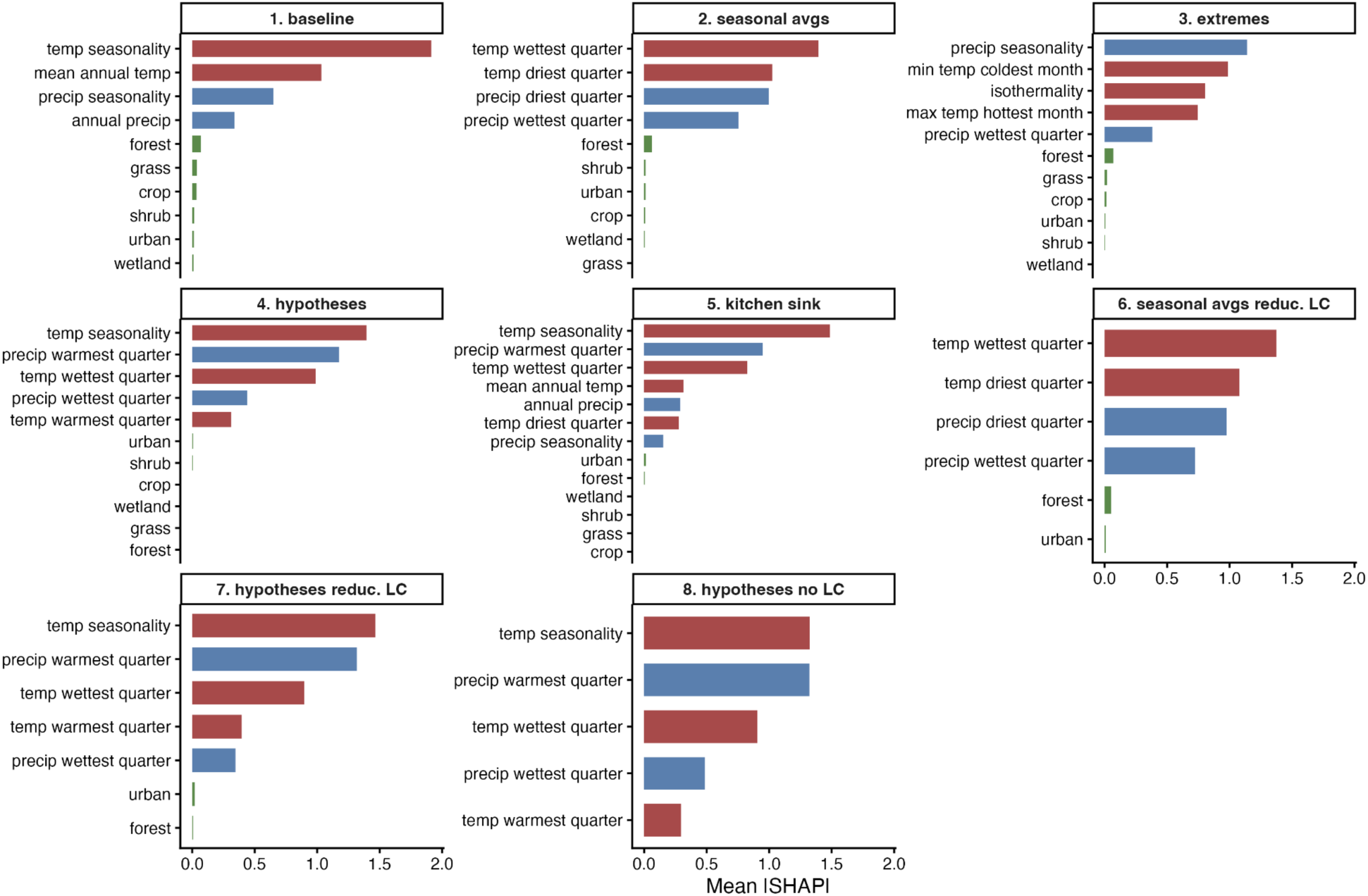
Variable importance (measured as the mean of the absolute value of the Shapley additive explanations [SHAP] score) for each model. Temperature variables are colored in red, precipitation variables in blue, and land cover variables in green. Panels are labeled by model number and nickname. Bioclimatic variables were more important than land cover variables in models that included both (models 1-7). Bioclimatic variables are labeled as: bio01 = "mean annual temp", bio02 = "mean DTR", bio03 = "isothermality", bio04 = "temp seasonality", bio05 = "max temp hottest month", bio06 = "min temp coldest month", bio07 = "annual temp range", bio08 = "temp wettest quarter", bio09 = "temp driest quarter", bio10 = "temp warmest quarter", bio11 = "temp coldest quarter", bio12 = "annual precip", bio13 = "precip wettest month", bio14 = "precip driest month", bio15 = "precip seasonality", bio16 = "precip wettest quarter", bio17 = "precip driest quarter", bio18 = "precip warmest quarter", bio19 = "precip coldest quarter". For model specifications, see Table 1.

Temperature played a large role in determining *Ae. sierrensis* habitat suitability. A temperature variable was ranked most important in all models except the extremes model (model 3), for which temperature was second behind precipitation; Fig. 3). Temperature seasonality (bio04) was the most important variable in every model in which it was included (models 1, 4, 5, 7, 8; Fig. 3), with the highest suitability at low temperature seasonality and suitability declining smoothly around 80 (in units of standard deviation of temperature in degrees Celsius *100; SHAP partial dependence plots [PDPs] for a subset of important bioclimatic variables shown in Fig. 4, all model PDPs shown in Figs. S8-S15). Mean temperature of the wettest quarter (bio08), representing average temperature during the season when *Ae. sierrensis* is actively developing, emerging, and reproducing, was the most important variable in the seasonal averages models (2, 6) and the third-most important variable in the models that included bio04 (hypotheses and kitchen sink models 4, 5, 7, 8; Figs. 4, S8-S15). Suitability peaked at average wettest-quarter temperatures of 0 – 10°C and declined sharply 10 – 20°C (Figs. 4, S8-S15). Temperature of the warmest quarter (bio10) showed a plateau of high suitability from 8 – 22°C but dropped steeply at higher temperatures (Figs. 4, S8-S15). For the extremes model (3), the minimum temperature of the coldest month (bio06) was the second most important variable (Fig. S10), suitability peaked around −5 – 5°C and declined steeply at both lower and higher temperatures.

**Figure 4.**
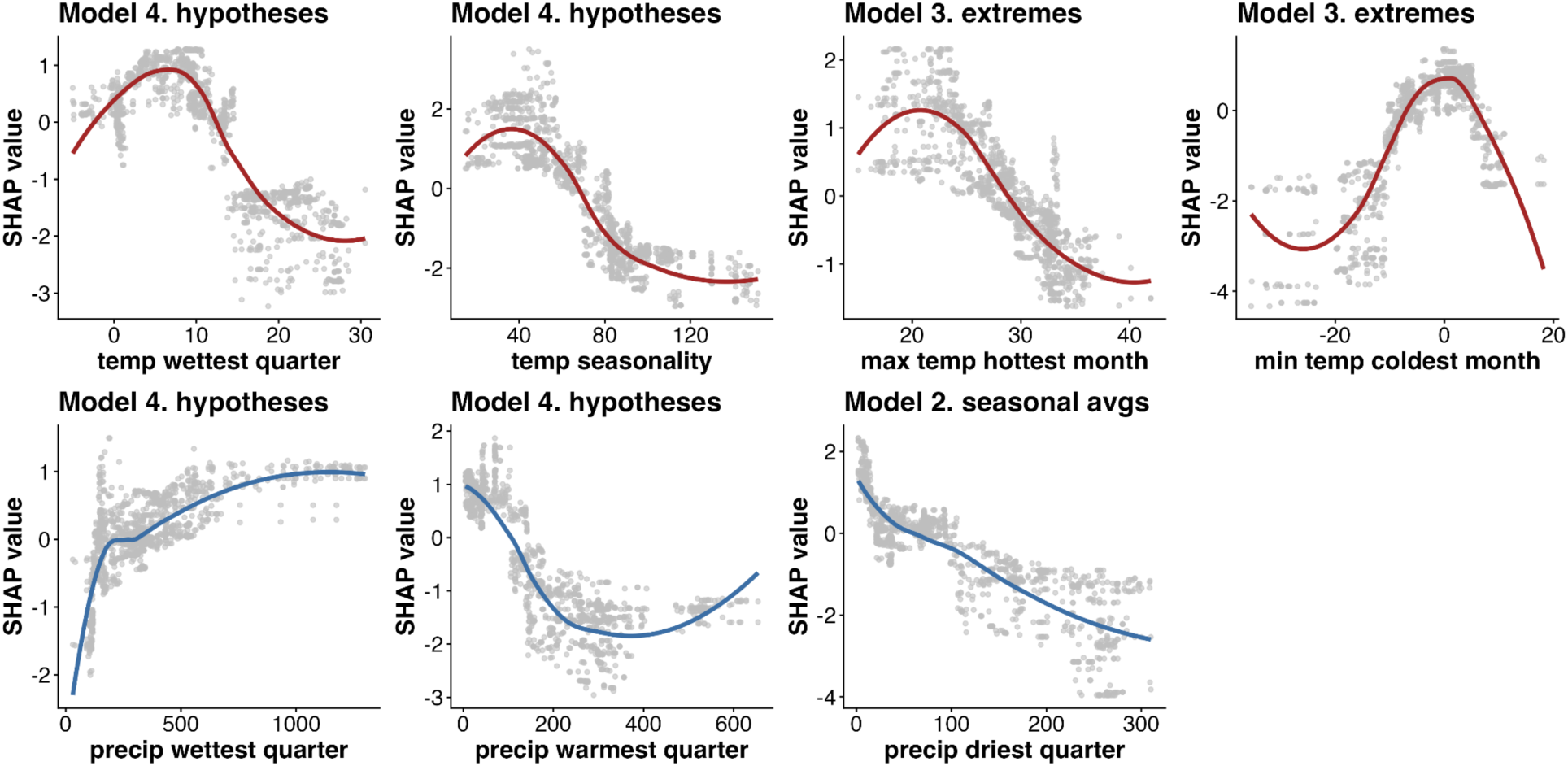
Nonlinear environmental responses: SHAP partial dependence plots show relationships between bioclimatic variables and habitat suitability for a subset of important variables in models 2, 3, and 4, which are characteristic of environmental responses across models. Plots in the top row illustrate unimodal relationships of habitat suitability to temperature variables, while those in the bottom row illustrate threshold-like relationships to precipitation variables. Panels are labeled by model number and nickname. Habitat suitability peaks with temperature of the wettest quarter (bio08) between 0 – 10°C, temperature seasonality (bio04) below 80, temperature of the warmest month (bio05) between 17 – 27°C, and temperature of the coldest month (bio06) between −5 – 5°C. Habitat suitability increases with precipitation in the wettest quarter (bio16) and decreases with precipitation in the warmest quarter (bio18), and driest quarter (bio17). Units are in °C (bio04, bio05, bio06, bio08), mm (bio16, bio17, bio18), or standard deviation*100 (bio04).

In addition to temperature, precipitation variables had high importance across all models. The most important precipitation variables were precipitation seasonality (bio15; baseline and extremes models 1, 3), precipitation of the driest quarter (bio17; seasonal averages models 2, 6), and precipitation of the warmest quarter (bio18; hypotheses and kitchen sink models 4, 5, 7, 8) (Fig. 3). Habitat suitability increased with greater precipitation seasonality, lower precipitation in the warmest quarter (sharply decreasing above 200 mm), and with lower precipitation in the driest quarter (Figs. 4, S8-S15). Other precipitation variables with moderate importance included precipitation of the wettest quarter (bio16; models 2, 3, 4, 6, 7, 8), which increased suitability sharply above ∼100 mm, and total annual precipitation (bio12; baseline model 1), which showed a nonlinearly increasing relationship with suitability (Figs. 4, S8-S15). Together, this indicates that environments with a pronounced dry season and sufficient rain in the wet season were important predictors of habitat suitability.

Land cover variables were always less important than all bioclimatic variables included in the models (Fig. 3). Where land cover variables showed any importance, forest was sometimes more important (baseline, seasonal average, and extremes models 1, 2, 3, and 6) as a positive predictor and urban was sometimes important (hypotheses and kitchen sink models 4, 5, and 7) as a negative predictor of suitability (Figs. 3-4, S8-S15). After forest and urban cover, the baseline model (1), which had the most sparse climate covariates, also showed some marginal importance of grassland, shrubland, wetland, and crop cover (Fig. 3).

In general, climate – suitability relationships tended to be nonlinear but smooth and gradual, rather than displaying either sharp thresholds or linear responses. Suitability responded unimodally to most temperature variables, with wet season temperature optima around 0 – 10°C average and −5 – 5°C minimum and 8 – 22°C non-active (warm) season temperatures, when *Ae. sierrensis* remains in a dormant dessication-resistant egg stage. Precipitation variables tended to show sharper thresholds, with suitability increasing steeply above 100 mm active-season precipitation and decreasing steeply above 100 mm warm-season precipitation (Figs. 4, S8-S15), delineating non-arid but seasonal, cool-season precipitation environments. In contrast to these thresholds, suitability linearly increased with precipitation seasonality and linearly decreased with precipitation of the driest quarter.

## Discussion

The geographic distribution of *Aedes sierrensis* is largely restricted to areas within a few hundred kilometers of the western coast of North America, from around the USA – Mexico border in the south through British Columbia and into southern Alaska in the north (Darsie and Ward 2005). Species distribution models captured this distribution with high confidence and accuracy (Figs. 2, S5-S7), generally making similar predictions despite using a variety of different bioclimatic and land cover variable predictors (Fig. S6, Table 1). Compared to other mosquito species that span a larger and more diverse geographic area (Fig. 1), *Ae. sierrensis* distribution may be particularly predictable. This may be because the species preference for tree hole breeding sites and its univoltine life cycle confine it to specific habitats with seasonal rainfall and trees (especially oak), and with limited competition with other mosquitoes using the same habitat (e.g., Kesavaraju et al. 2014), as well as because it occupies its native range and is not expanding into new regions.

The models predicted high-confidence regions of the distribution along the Pacific coast and interior regions, running from northern Baja California in the south to British Columbia and southern Alaska in the north (Fig. 2). In addition, models predicted intermediate to high suitability with lower confidence (greater variation among models) in inland regions of the Intermountain West, USA, western Mexico (Sierra Madre Occidental), and small pockets of southern Mexico (in the highlands near Mexico City) (Fig. S6). The models consistently predicted lower suitability in the California Central Valley, USA and southern Baja California, Mexico, areas adjacent to the species range where summer temperatures and aridity limit habitat suitability.

Because bioclimatic variables can be highly correlated (Fig. S2), multiple combinations of variables can describe similar regions of environmental space (i.e., temperature and precipitation over different time ranges). To aid model interpretability and reduce overfitting, I constructed a series of models based on narratives characterizing climate variation in different ways, each including a reduced set of 2-3 temperature variables, 2-3 precipitation variables, and 0, 2, or 6 land cover variables (Table 1). These narratives included a baseline climate model that included annual average temperature and precipitation and their seasonal variation (model 1), a model based on seasonal averages (model 2), a model capturing variability and extremes (model 3), a model incorporating biological hypotheses about how active (wet) season and inactive (hot) season temperature and precipitation affect mosquito performance (model 4), a ‘kitchen sink’ model including these and other variables (model 5), and variants on models 2 and 4 that removed some or all land cover variables (models 6-8; Table 1). Because these variables capture overlapping variation, flexible XGBoost models could arrive at similar performance, prediction maps, environmental variable importance, and environment – suitability relationships (comparing different but related bioclimatic variables) using different predictor sets. While this consistency increases confidence in model predictions, it also limits more precise mechanistic interpretation of which climate factors directly limit the species distribution. Synthesizing across bioclimatic variables and model specifications, models support the hypothesis that *Ae. sierrensis* is constrained to climates with low to moderate, seasonally stable temperatures and moderate, seasonal rainfall concentrated in the cool season, consistent with Marine/Oceanic West Coast, Mediterranean, and other dry-summer climates of western North America. These climates often coincide with forested environments (Fig. S1).

Despite overall model agreement throughout most of the predicted range, model comparison highlights several areas of uncertainty in *Ae. sierrensis* habitat suitability, typically in regions with limited mosquito sampling (for both the focal and background species). The areas of highest uncertainty (i.e., standard deviation in predicted suitability among models; Fig. 2B) coincide with regions with limited to no occurrence or background points used for model fitting (Fig. 1), including the Intermountain West, central-west Mexico, the southeastern USA, and Canada between the Coast and Rocky Mountains. While I purposely searched for mosquito records (i.e., background points) spanning the entire geography of Canada, USA, and Mexico (Table S1), the 2:1 ratio of background to occurrence points—imposed to ensure balance and to avoid overfitting on pseudo-absences—constrained the ability to fully cover the multidimensional environmental space. Ultimately, the availability of spatially independent *Ae. sierrensis* occurrence points is the key constraint on how comprehensively the background points can represent geographic and environmental space. Field mosquito sampling could help to resolve uncertainty in the suitability in central Mexico, where mosquito observations are infrequent in publicly available databases like GBIF and other US-based sources like VectorSurv.

Temperature variables were typically the most important for delineating *Ae. sierrensis* habitat suitability, closely followed by precipitation. SDMs estimated unimodal relationships between temperature variables and habitat suitability, consistent with theory and laboratory experimental work on organismal trait thermal performance (Mordecai et al. 2019), suggesting that individual-level physiological constraints shape species distributions. However, SDM-estimated temperature responses trended cooler than those previously measured in a laboratory experiment. For 10 populations of *Ae. sierrensis* collected from across a 1,200 km gradient and reared in a common garden laboratory experiment at a range of constant temperatures, larval and pupal survival peaked from 10 – 25°C, adult survival peaked at 15 – 17°C, and immature development rate peaked at 26 – 27°C (Couper et al. 2024a). The experimental lower thermal limits for survival at all life stages ranged from 0 – 2°C, while upper thermal limits ranged from 32 – 34°C (Couper et al. 2024a). By contrast, based on the SDMs, occurrence is constrained to average temperatures 5 – 10°C lower than these experimental optima: average wet-season temperatures of 0 – 10°C, minimum temperatures of the coolest month of −5 – 5°C, and summer temperatures of 8 – 22°C. In some cooler parts of the range where the wettest quarter is a snowy winter, this may indicate a mismatch between the actual larval activity season (spring, when snowmelt begins to fill tree holes) and the chosen temperature variable. This comparison of lab and SDM results indicates that while *Ae. sierrensis* can tolerate and even thrive at warmer temperatures in a controlled laboratory environment, it is limited to cooler regions in the field. This is consistent with findings of a recent study that compared thermal limit estimates from SDMs to those from mechanistic laboratory experiments for seven major mosquito vector species, which found a high correlation (r = 0.869 across species) between the thermal minimum measured by SDM versus lab experiment, but with systematically ∼5°C lower thermal limits from SDMs for *Ae. aegypti* and *Ae. albopictus* (Athni et al. 2024).

Results showing correlated but lower thermal optima in the field than measured at constant temperatures in the lab are consistent with ectotherm physiology empirical work and theory demonstrating the importance of thermal safety margins, in which organisms typically live in environments where average temperatures are below their performance optima to buffer against the harmful effects of temperature variation and heat extremes (Kingsolver 2009, Clusella-Trullas et al. 2021, Buckley et al. 2022, Couper et al. 2024b). By contrast, peak habitat suitability for *Ae. sierrensis* coincides with temperatures that maximize infection with the parasite *Lambornella clarki* in both the laboratory and field (Farner et al. 2026), suggesting that while temperature-dependent parasitism may limit individual fitness, abundance, and population dynamics, it does not constrain the distribution of *Ae. sierrensis*. Taken together, experimental work (Couper et al. 2024a) suggests that pupal, and to a lesser extent larval, survival constrain the lower thermal limits of the *Ae. sierrensis* distribution, while larval and adult survival constrain its upper thermal limits, and parasitism may limit abundance but not the species range (Farner et al. 2026).

Compared to climate, land cover variables were consistently less important for predicting habitat suitability. Within models that included land cover variables, they were consistently ranked below all bioclimatic variables based on SHAP scores and gain (Figs. 3, S7), and a model that excluded land cover produced similar predictive accuracy, suitability map, variable importance, and environmental responses as models that included land cover variables (i.e., comparing model 8 to models 4 and 6). Because climate variables directly determine vegetation classes such as forest, shrubland, grassland, and wetland, this may indicate that climate variables subsume most of the variation in land cover but explain more variation in habitat suitability because climate also directly affects the mosquito. An additional potential explanation is that the land cover variables themselves are not highly predictive of *Ae. sierrensis* occurrence. The land cover variables included in the model were modal land cover classes within a 1-km circular buffer, chosen to represent the area immediately surrounding the sampling site; it is possible that land cover variables calculated at either larger spatial scales to capture the broader regional environment or at hyper-local scales to capture the precise environment of the collection site might have more predictive power. Moreover, more specific habitat indicators such as cover of oaks could be stronger predictors of *Ae. sierrensis* occurrence. However, the high performance of model 8, which excluded land cover variables (i.e., 0.94 out-of-sample AUC) shows that climate variables leave little variation for land cover to potentially explain.

A growing body of research uses SDMs to understand the current drivers of mosquito species ranges and to project their future changes under global change scenarios, consistently finding that temperature is among the most important predictors (Lippi et al. 2023). A systematic review identified 204 studies from the last 20 years that developed SDMs for mosquitoes, covering 138 mosquito species but primarily focusing on major disease vector species like *Ae. aegypti*, *Ae. albopictus*, *Cx. pipiens*, and *Anopheles gambiae*, and primarily using MaxEnt methods (only 6.9% of studies used regression tree methods like those used here) (Lippi et al. 2023). As in this work, temperature was the most important predictor of habitat suitability in 54.9% of studies, followed by precipitation (42.6% of studies), and land cover and land use (31.4% of studies) (Lippi et al. 2023). For example, a prominent SDM for the globally invasive mosquito vectors *Ae. aegypti* and *Ae. albopictus* found that temperature was the primary predictor of habitat suitability, while precipitation followed by vegetation explained most of the rest of the variation, with little importance of urban cover (Kraemer et al. 2015). By contrast, several models of *Culex* spp. vectors of West Nile virus have found high importance of built environment, human population density, and land cover, as well as temperature, precipitation, and specific humidity (Larson et al. 2010, Gorris et al. 2021, Rhodes et al. 2023). Taken together, these studies highlight that the thermal biology of mosquitoes (Mordecai et al. 2019), combined with their sensitivity to relative humidity (Brown et al. 2023, Huxley et al. 2026) and water accumulation, either through precipitation or human water storage (depending on mosquito oviposition habitat preference) (Lowe et al. 2021) acting at individual and population scales determine predictable species-level habitat constraints.

While SDMs can be powerful tools for generating, and sometimes testing, hypotheses about the ecological determinants of species ranges, recent literature has recommended caution in using them for applied management decisions, suggesting best practices for modeling and reproducibility (Araújo et al. 2019, Feng et al. 2019, Zurell et al. 2020, Barker and MacIsaac 2022). A review of 127 mosquito SDMs published between 1998-2020 found that many used one or more practices that are considered unacceptable for application to biodiversity assessments or mosquito control decisions (Barker and MacIsaac 2022). In this work, I made efforts to apply the recommended best practices for SDM modeling and reproducibility (Araújo et al. 2019, Feng et al. 2019, Zurell et al. 2020, Barker and MacIsaac 2022), including through thoughtful selection of occurrence and background points, hypothesis-driven environmental covariates at relevant spatial and temporal scales, model training and testing with spatial block cross-validation to avoid overfitting and spatial autocorrelation, multi-model comparison, and ensemble modeling. However, modeling methods could be improved to approach gold standards by conducting new primary field sampling to directly test predictions (particularly in undersampled geographic areas), by enhancing the temporal match between environmental predictors and occurrence points, by improving methods for propagating uncertainty from response and predictor variables through to model fitting, by applying multiple algorithmic approaches (e.g., MaxEnt, random forest), and by directly testing environmental responses with controlled experiments. In particular, mosquito sampling in tree-covered areas of the Intermountain West, USA, and highland western and central Mexico would help to resolve uncertainty in the distributional limits of *Ae. sierrensis*. These remain useful directions for future research.

This work focused on understanding the environmental drivers of the current distribution of *Ae. sierrensis*, but it also has implications for the species’ response to climate change. The models identified constraints on long-term seasonal or monthly average temperatures that were 5 – 10°C cooler than laboratory-measured constant temperature responses. These cooler realized thermal limits imply that the species may be sensitive to warming temperature as warmer winters and summers potentially reduce suitability at the southern range extent while expanding suitability further north into coastal Alaska. Precipitation changes have a more uncertain impact both because they are more uncertain in climate model projections and because they have sharper thresholds in the SDMs. Climate variables synergistically affect *Ae. sierrensis* habitat suitability: for example, changes in the duration, timing, and intensity of the rainy season in coastal and Mediterranean climates affect when and for how long the mosquito is active at immature and adult stages, in turn affecting the range of temperatures it experiences in these life stages. Biotic interactions may further mediate the species’ response to climate variation and change: previous field work found that while parasitism consistently peaked in the center of the species range with wet-season temperatures of 10°C across years, parasitism was more common in a cooler, wetter year than in a warmer, drier year (Farner et al. 2026). Finally, changes in the availability of tree hole habitat, e.g., through climate-driven range shifts of oaks (Kueppers et al. 2005), could indirectly affect the extent of climatically suitable habitat that can be occupied by *Ae. sierrensis*. Taken together, this suggests that general predictions like a climate-driven northward range shift may be reasonable while more specific predictions may depend on interactions between precipitation, temperature, biotic interactions, and evolutionary and behavioral adaptation.

## Conclusions

Understanding fundamental environmental constraints and the realized range in which species occur has long been a central goal in ecology (Elton 1927, Hutchinson 1957). Recent advances in large-scale biodiversity and environmental data, along with flexible computational methods from machine learning, have accelerated the pace of discovery. At the same time, mechanistic understanding of ecological constraints, and their implications for global change, requires integrating field observations, laboratory experiments, large-scale environmental data, and modeling. Here, I develop the first species distribution models for *Ae. sierrensis*, a widespread mosquito that is ecologically important throughout western North America, a nuisance mosquito, and a vector of dog heartworm. Habitat suitability was highly predictable from long-term averages of seasonal temperature and precipitation. Climatic responses were consistently nonlinear: temperature variables tended to have smooth, hump-shaped relationships with habitat suitability while precipitation variables tended to have sharper thresholds. These responses are consistent with mechanistic evidence of the importance of temperature in the laboratory, but differ in their absolute thermal optima and limits. Because different bioclimatic variables capture overlapping aspects of the environment, models including different variable sets produced similar performance and predictions. As a result, interpreting the causal variables underlying species distributions requires integrating observational evidence with mechanistic knowledge of species life cycles and responses to environmental variation, including through laboratory and field validation, returning to the centuries-old grounding of ecology in natural history.

## Acknowledgements

Many Mordecai lab members provided generous feedback that substantially improved this work. Kelsey Lyberger provided the photos in Figure 1. Caroline Glidden, Aly Singleton, Lisa Couper, and Sam Sambado provided modeling guidance and example code. Isabel Delwel was my learning partner as we learned about species distribution modeling together. Otto Seppälä first introduced me to tree hole mosquitoes and their parasites, and Jan Washburn’s pioneering research and personal guidance supported our first forays into research in this fascinating system. This work was funded by the National Institutes of Health (R35GM13439).

## Author contributions

Erin Mordecai conducted all data extraction, analyses, visualization, and manuscript writing.

## References

Araújo, M. B., R. P. Anderson, A. Márcia Barbosa, C. M. Beale, C. F. Dormann, R. Early, R. A. Garcia, A. Guisan, L. Maiorano, B. Naimi, R. B. O’Hara, N. E. Zimmermann, and C. Rahbek. 2019. Standards for distribution models in biodiversity assessments. Science Advances 5:eaat4858.

Athni, T. S., M. L. Childs, C. K. Glidden, and E. A. Mordecai. 2024. Temperature dependence of mosquitoes: Comparing mechanistic and machine learning approaches. PLOS Neglected Tropical Diseases 18:e0012488.

Barker, J. R., and H. J. MacIsaac. 2022. Species distribution models applied to mosquitoes: Use, quality assessment, and recommendations for best practice. Ecological Modelling 472:110073.

Barrett, T., M. Dowle, A. Srinivasan, J. Gorecki, M. Chirico, T. Hocking, B. Schwendinger, I. Krylov, P. Stetsenko, T. Short, S. Lianoglou, E. Antonyan, M. Bonsch, H. Parsonage, S. Ritchie, K. Ren, X. Tan, R. Saporta, O. Seiskari, X. Dong, M. Lang, W. Iwasaki, S. Wenchel, K. Broman, T. Schmidt, D. Arenburg, E. Smith, F. Cocquemas, M. Gomez, P. Chataignon, N. Blaser, D. Selivanov, A. Riabushenko, C. Lee, D. Groves, D. Possenriede, F. Parages, D. Toth, M. Yaramaz-David, A. Perumal, J. Sams, M. Morgan, M. Quinn, @javrucebo (GitHub user), M. Halperin, R. Storey, M. Saraswat, M. Jacob, M. Schubmehl, D. Vaughan, L. Silvestri, J. Hester, A. Damico, S. Freundt, D. Simons, E. S. de Andrade, C. Miller, J. P. Meldgaard, V. Tlapak, K. Ushey, D. Eddelbuettel, T. Fischetti, O. Shilon, V. Khotilovich, H. Wickham, B. Becker, K. Haynes, B. C. Kamgang, O. Delmarcell, J. O’Brien, D. de Mezquita, M. Czekanski, D. Shemetov, N. Jha, J. Wu, I. Giné-Vázquez, A. Chetia, D. Amoakohene, A. Feliz, M. Young, M. Seeto, P. Grosjean, V. Runge, C. Wia, E. Maigné, V. Rocher, V. Lulla, A. Sluga, B. Evans, R. Bruner, @badasahog (GitHub user), V. Thakur, M. Kumar, I. Czeller, M. Das, and T. Thammisetty. 2026, May 6. data.table: Extension of “data.frame.”

Bengtsson, H., and R Core Team. 2026, February 20. future.apply: Apply Function to Elements in Parallel using Futures.

Bisong, E. 2019. Google Colaboratory. Pages 59–64 in E. Bisong, editor. Building Machine Learning and Deep Learning Models on Google Cloud Platform: A Comprehensive Guide for Beginners. Apress, Berkeley, CA.

Brown, J. J., M. Pascual, M. C. Wimberly, L. R. Johnson, and C. C. Murdock. 2023. Humidity – The overlooked variable in the thermal biology of mosquito-borne disease. Ecology Letters 26:1029– 1049.

Buckley, L. B., R. B. Huey, and J. G. Kingsolver. 2022. Asymmetry of thermal sensitivity and the thermal risk of climate change. Global Ecology and Biogeography 31:2231–2244.

Campitelli, E. 2025, June 20. ggnewscale: Multiple Fill and Colour Scales in “ggplot2.”

Chamberlain, S., D. Oldoni, V. Barve, P. Desmet, L. Geffert, D. Mcglinn, K. Ram, rOpenSci, and J. Waller. 2026, March 20. rgbif: Interface to the Global Biodiversity Information Facility API.

Chen, T., T. He, M. Benesty, V. Khotilovich, Y. Tang, H. Cho, K. Chen, R. Mitchell, I. Cano, T. Zhou, M. Li, J. Xie, M. Lin, Y. Geng, Y. Li, J. Yuan, D. Cortes, and Xgb. contributors (base Xgb. implementation). 2026, March 18. xgboost: Extreme Gradient Boosting.

Clusella-Trullas, S., R. A. Garcia, J. S. Terblanche, and A. A. Hoffmann. 2021. How useful are thermal vulnerability indices? Trends in Ecology & Evolution 36:1000–1010.

Commission for Environmental Cooperation (CEC), Canada Centre for Remote Sensing (CCRS), U.S. Geological Survey (USGS), Comisión Nacional para el Conocimiento y Uso de la Biodiversidad (CONABIO), Comisión Nacional Forestal (CONAFOR), and Instituto Nacional de Estadística y Geografía (INEGI). 2024. North American Environmental Atlas - Land Cover 2020 30m. Raster digital data [30-m], Earth Engine Data Catalog.

Couper, L. I., T. O. Dodge, J. A. Hemker, B. Y. Kim, M. Exposito-Alonso, R. B. Brem, E. A. Mordecai, and M. C. Bitter. 2025. Evolutionary adaptation under climate change: Aedes sp. demonstrates potential to adapt to warming. Proceedings of the National Academy of Sciences 122:e2418199122.

Couper, L. I., J. E. Farner, K. P. Lyberger, A. S. Lee, and E. A. Mordecai. 2024a. Mosquito thermal tolerance is remarkably constrained across a large climatic range. Proceedings of the Royal Society B: Biological Sciences 291:20232457.

Couper, L. I., and E. A. Mordecai. 2022. Ecological drivers of dog heartworm transmission in California. Parasites & Vectors 15:388.

Couper, L. I., D. U. Nalukwago, K. P. Lyberger, J. E. Farner, and E. A. Mordecai. 2024b. How Much Warming Can Mosquito Vectors Tolerate? Global Change Biology 30:e17610.

Darsie, R. F., and R. A. Ward. 2005. Identification and Geographical Distribution of the Mosquitoes of North America, North of Mexico. University of Florida Press, Gainesville, Florida, USA.

Elith, J., C. H. Graham, R. P. Anderson, M. Dudík, S. Ferrier, A. Guisan, R. J. Hijmans, F. Huettmann, J. R. Leathwick, A. Lehmann, J. Li, L. G. Lohmann, B. A. Loiselle, G. Manion, C. Moritz, M. Nakamura, Y. Nakazawa, J. M. M. Overton, A. T. Peterson, S. J. Phillips, K. Richardson, R. Scachetti-Pereira, R. E. Schapire, J. Soberón, S. Williams, M. S. Wisz, and N. E. Zimmermann. 2006. Novel methods improve prediction of species’ distributions from occurrence data. Ecography 29:129–151.

Elith, J., J. R. Leathwick, and T. Hastie. 2008. A working guide to boosted regression trees. Journal of Animal Ecology 77:802–813.

Elton, C. S. 1927. Animal Ecology. University of Chicago Press.

Farner, J. E., K. P. Lyberger, L. I. Couper, M. Cruz-Loya, and E. A. Mordecai. 2026. Nonlinear effects of temperature on mosquito parasite infection across a large geographic climate gradient. Ecology 107:e70307.

Feng, X., D. S. Park, C. Walker, A. T. Peterson, C. Merow, and M. Papeş. 2019. A checklist for maximizing reproducibility of ecological niche models. Nature Ecology & Evolution 3:1382– 1395.

Foley, J. A., R. DeFries, G. P. Asner, C. Barford, G. Bonan, S. R. Carpenter, F. S. Chapin, M. T. Coe, G. C. Daily, H. K. Gibbs, J. H. Helkowski, T. Holloway, E. A. Howard, C. J. Kucharik, C. Monfreda, J. A. Patz, I. C. Prentice, N. Ramankutty, and P. K. Snyder. 2005. Global Consequences of Land Use. Science 309:570–574.

Frick, H., F. Chow, M. Kuhn, M. Mahoney, J. Silge, H. Wickham, P. Software, PBC [cph, and fnd. 2026, January 30. rsample: General Resampling Infrastructure.

Gorelick, N., M. Hancher, M. Dixon, S. Ilyushchenko, D. Thau, and R. Moore. 2017. Google Earth Engine: Planetary-scale geospatial analysis for everyone. Remote Sensing of Environment 202:18–27.

Gorman, B. 2018, May 12. mltools: Machine Learning Tools.

Gorris, M. E., A. W. Bartlow, S. D. Temple, D. Romero-Alvarez, D. P. Shutt, J. M. Fair, K. A. Kaufeld, S. Y. Del Valle, and C. A. Manore. 2021. Updated distribution maps of predominant Culex mosquitoes across the Americas. Parasites & Vectors 14:547.

Greenwell, B. 2026a, July 5. bgreenwell/vip. R.

Greenwell, B. M. 2026b, January 23. pdp: Partial Dependence Plots.

Hijmans, R. J., A. Brown, M. Barbosa, K. Dyba, R. Bivand, M. Chirico, E. Cordano, E. Pebesma, B. Rowlingson, and M. D. Sumner. 2026, June 19. terra: Spatial Data Analysis.

Hijmans, R. J., S. E. Cameron, J. L. Parra, P. G. Jones, and A. Jarvis. 2005. Very high resolution interpolated climate surfaces for global land areas. International Journal of Climatology 25:1965– 1978.

Hijmans, R. J., J. van Etten, M. Sumner, J. Cheng, D. Baston, A. Bevan, R. Bivand, L. Busetto, M. Canty, B. Fasoli, D. Forrest, A. Ghosh, D. Golicher, J. Gray, J. A. Greenberg, P. Hiemstra, K. Hingee, A. Ilich, I. for M. A. Geosciences, C. Karney, M. Mattiuzzi, S. Mosher, B. Naimi, J. Nowosad, E. Pebesma, O. P. Lamigueiro, E. B. Racine, B. Rowlingson, A. Shortridge, B. Venables, and R. Wueest. 2025, March 28. raster: Geographic Data Analysis and Modeling.

Hijmans, R. J., S. Phillips, J. Leathwick, and J. Elith. 2024, November 25. dismo: Species Distribution Modeling.

Hutchinson, G. E. 1957. Concluding Remarks. Cold Spring Harbor Symposia on Quantitative Biology 22:415–427.

Huxley, P. J., J. J. Brown, B. St. Laurent, B. Johnson, O. Y. Cheung, A. Asamoah, B. D. Hollingsworth, E. R. Bump, M. C. Wimberly, M. Pascual, L. R. Johnson, and C. C. Murdock. 2026. Beyond Temperature: Relative Humidity Systematically Shifts Juvenile Thermal Performance and Projected Population Growth in a Malaria Vector. Ecology Letters 29:e70416.

Ismail, S., J. Farner, L. Couper, E. Mordecai, and K. Lyberger. 2023. Temperature and intraspecific variation affect host–parasite interactions. Oecologia.

Kesavaraju, B., P. T. Leisnham, S. Keane, N. Delisi, and R. Pozatti. 2014. Interspecific Competition between Aedes albopictus and A. sierrensis: Potential for Competitive Displacement in the Western United States. PLOS ONE 9:e89698.

Kingsolver, J. G. 2009. The Well-Temperatured Biologist: (American Society of Naturalists Presidential Address). The American Naturalist 174:755–768.

Kraemer, M. U., M. E. Sinka, K. A. Duda, A. Q. Mylne, F. M. Shearer, C. M. Barker, C. G. Moore, R. G. Carvalho, G. E. Coelho, W. Van Bortel, G. Hendrickx, F. Schaffner, I. R. Elyazar, H.-J. Teng, O. J. Brady, J. P. Messina, D. M. Pigott, T. W. Scott, D. L. Smith, G. W. Wint, N. Golding, and S. I. Hay. 2015. The global distribution of the arbovirus vectors Aedes aegypti and Ae. albopictus. eLife 4.

Kueppers, L. M., M. A. Snyder, L. C. Sloan, E. S. Zavaleta, and B. Fulfrost. 2005. Modeled regional climate change and California endemic oak ranges. Proceedings of the National Academy of Sciences 102:16281–16286.

Kuhn, M. 2008. Building Predictive Models in R Using the caret Package. Journal of Statistical Software 28:1–26.

Larson, S. R., J. P. Degroot, L. C. Bartholomay, and R. Sugumaran. 2010. Ecological niche modeling of potential West Nile virus vector mosquito species in Iowa. Journal of Insect Science 10:110.

Ledesma, N., and L. Harrington. 2011. Mosquito Vectors of Dog Heartworm in the United States: Vector Status and Factors Influencing Transmission Efficiency. Topics in Companion Animal Medicine 26:178–185.

Lippi, C. A., S. J. Mundis, R. Sippy, J. M. Flenniken, A. Chaudhary, G. Hecht, C. J. Carlson, and S. J. Ryan. 2023. Trends in mosquito species distribution modeling: insights for vector surveillance and disease control. Parasites & Vectors 16:302.

Liu, Y., A. Just, and M. Mayer. 2025, December 13. SHAPforxgboost: SHAP Plots for “XGBoost.”

Lowe, R., S. A. Lee, K. M. O’Reilly, O. J. Brady, L. Bastos, G. Carrasco-Escobar, R. de Castro Catão, F. J. Colón-González, C. Barcellos, M. S. Carvalho, M. Blangiardo, H. Rue, and A. Gasparrini. 2021. Combined effects of hydrometeorological hazards and urbanisation on dengue risk in Brazil: a spatiotemporal modelling study. The Lancet Planetary Health 5:e209–e219.

Ludlow, C. S. 1905. Mosquito Notes.-No. 3. The Canadian Entomologist 37:133–135.

Lyberger, K., J. Farner, L. Couper, and E. A. Mordecai. 2024. A Mosquito Parasite Is Locally Adapted to Its Host but Not Temperature. The American Naturalist 204:121–132.

Mahoney, M., J. Silge, P. Software, and PBC. 2025, December 2. spatialsample: Spatial Resampling Infrastructure.

Massicotte, P., A. South, and K. Hufkens. 2026, January 19. rnaturalearth: World Map Data from Natural Earth.

Matthiopoulos, J. 2022. Defining, estimating, and understanding the fundamental niches of complex animals in heterogeneous environments. Ecological Monographs 92:e1545.

Mordecai, E. A., J. M. Caldwell, M. K. Grossman, C. A. Lippi, L. R. Johnson, M. Neira, J. R. Rohr, S. J. Ryan, V. Savage, M. S. Shocket, R. Sippy, A. M. S. Ibarra, M. B. Thomas, and O. Villena. 2019. Thermal biology of mosquito-borne disease. Ecology Letters 22:1690–1708.

Neuwirth, E. 2022, April 3.RColorBrewer: ColorBrewer Palettes.

Pebesma, E., R. Bivand, E. Racine, M. Sumner, I. Cook, T. Keitt, R. Lovelace, H. Wickham, J. Ooms, K. Müller, T. L. Pedersen, D. Baston, D. Dunnington, and A. Courtiol. 2026, May 6. sf: Simple Features for R.

Pedersen, T. L. 2025, August 25. patchwork: The Composer of Plots.

Peterson, B. G., P. Carl, K. Boudt, R. Bennett, J. Ulrich, E. Zivot, D. Cornilly, E. Hung, M. Lestel, K. Balkissoon, D. Wuertz, A. A. Christidis, R. D. Martin, Z. Z. Zhou, J. M. Shea, D. Jain, and T. Daniyarov. 2026, April 11. PerformanceAnalytics: Econometric Tools for Performance and Risk Analysis.

Pulliam, H. r. 2000. On the relationship between niche and distribution. Ecology Letters 3:349–361.

Queiroz, G. D., C. Fay, E. Hvitfeldt, O. Keyes, K. Misra, T. Mastny, J. Erickson, D. Robinson, J. Silge [aut, and cre. 2025, July 25. tidytext: Text Mining using “dplyr”, “ggplot2”, and Other Tidy Tools.

Rhodes, C. G., L. F. Chaves, L. R. Bergmann, and G. L. Hamer. 2023. Ensemble species distribution modeling of Culex tarsalis (Diptera: Culicidae) in the continental United States. Journal of Medical Entomology 60:664–679.

Robin, X., N. Turck, A. Hainard, N. Tiberti, F. Lisacek, J.-C. Sanchez, M. Müller, S. S. (Fast D. code), M. D. (Hand & T. Multiclass), and Z. B. (DeLong paired test CI). 2025, July 31. pROC: Display and Analyze ROC Curves.

Rockström, J., W. Steffen, K. Noone, Å. Persson, F. S. Chapin, E. F. Lambin, T. M. Lenton, M. Scheffer, C. Folke, H. J. Schellnhuber, B. Nykvist, C. A. de Wit, T. Hughes, S. van der Leeuw, H. Rodhe, S. Sörlin, P. K. Snyder, R. Costanza, U. Svedin, M. Falkenmark, L. Karlberg, R. W. Corell, V. J. Fabry, J. Hansen, B. Walker, D. Liverman, K. Richardson, P. Crutzen, and J. A. Foley. 2009. A safe operating space for humanity. Nature 461:472–475.

Singleton, A. L., C. K. Glidden, A. J. Chamberlin, R. Tuan, R. G. S. Palasio, A. Pinter, R. L. Caldeira, C. L. F. Mendonça, O. S. Carvalho, M. V. Monteiro, T. S. Athni, S. H. Sokolow, E. A. Mordecai, and G. A. D. Leo. 2024. Species distribution modeling for disease ecology: A multi-scale case study for schistosomiasis host snails in Brazil. PLOS Global Public Health 4:e0002224.

Taiyun. 2026, June 22. taiyun/corrplot. R.

Valavi, R., J. Elith, J. Lahoz-Monfort, I. Flint, and G. Guillera-Arroita. 2025, August 21. blockCV: Spatial and Environmental Blocking for K-Fold and LOO Cross-Validation.

Villeneuve, C.-A., J. Snyman, L. P. Snyman, G. G. Gouin, E. Jenkins, V. Martinez, T. Hobman, A. Kumar, I. Dusfour, N. Lecomte, and P. A. Leighton. 2025. Expanding knowledge of mosquito (Diptera: Culicidae) and California serogroup viruses distributions in the North American Arctic. Journal of Medical Entomology 62:1590–1598.

Washburn, J. O., J. R. Anderson, and D. R. Mercer. 1989. Emergence Characteristics of Aedes sierrensis (Diptera: Culicidae) from California Treeholes with Particular Reference to Parasite Loads. Journal of Medical Entomology 26:173–182.

Washburn, J. O., M. E. Gross, D. R. Mercer, and J. R. Anderson. 1988. Predator-induced trophic shift of a free-living ciliate: parasitism of mosquito larvae by their prey. Science 240:1193–1195.

Washburn, J. O., D. R. Mercer, and Anderson. 1991. Regulatory role of parasites: impact on host population shifts with resource availability. Science 253:185–188.

Wickham, H., W. Chang, L. Henry, T. L. Pedersen, K. Takahashi, C. Wilke, K. Woo, H. Yutani, D. Dunnington, T. van den Brand, Posit, and PBC. 2026a, April 22. ggplot2: Create Elegant Data Visualisations Using the Grammar of Graphics.

Wickham, H., R. François, L. Henry, K. Müller, D. Vaughan, P. Software, and PBC. 2026b, April 3. dplyr: A Grammar of Data Manipulation.

Wickham, H., L. Henry, P. Software, PBC [cph, and fnd. 2026c, April 10. purrr: Functional Programming Tools.

Wickham, H., and RStudio. 2023, February 22. tidyverse: Easily Install and Load the “Tidyverse.”

Wickham, H., P. Software, PBC [cph, and fnd. 2025a, September 25. forcats: Tools for Working with Categorical Variables (Factors).

Wickham, H., D. Vaughan, M. Girlich, K. Ushey, P. Software, and PBC. 2025b, December 19. tidyr: Tidy Messy Data.

Wood, S. 2025, November 7. mgcv: Mixed GAM Computation Vehicle with Automatic Smoothness Estimation.

Yan, Y. 2025, November 14. rBayesianOptimization: Bayesian Optimization of Hyperparameters.

Yee, W. L., and J. R. Anderson. 1995. Tethered Flight Capabilities and Survival of Lambornella clarki-Infected, Blood-Fed, and Gravid Aedes sierrensis (Diptera: Culicidae). Journal of Medical Entomology 32:153–160.

Zurell, D., J. Franklin, C. König, P. J. Bouchet, C. F. Dormann, J. Elith, G. Fandos, X. Feng, G. Guillera-Arroita, A. Guisan, J. J. Lahoz-Monfort, P. J. Leitão, D. S. Park, A. T. Peterson, G. Rapacciuolo, D. R. Schmatz, B. Schröder, J. M. Serra-Diaz, W. Thuiller, K. L. Yates, N. E. Zimmermann, and C. Merow. 2020. A standard protocol for reporting species distribution models. Ecography 43:1261–1277

