## Supplementary Information for "Nonlinear responses to temperature and precipitation shape the distribution of *Aedes sierrensis* in North America"

Contents:
Appendix S1: Supplementary Figures S1-S15
Appendix S2: Supplementary Tables S1-S2
Appendix S3: ODMAP Protocol

### Appendix S1. Supplementary Figures

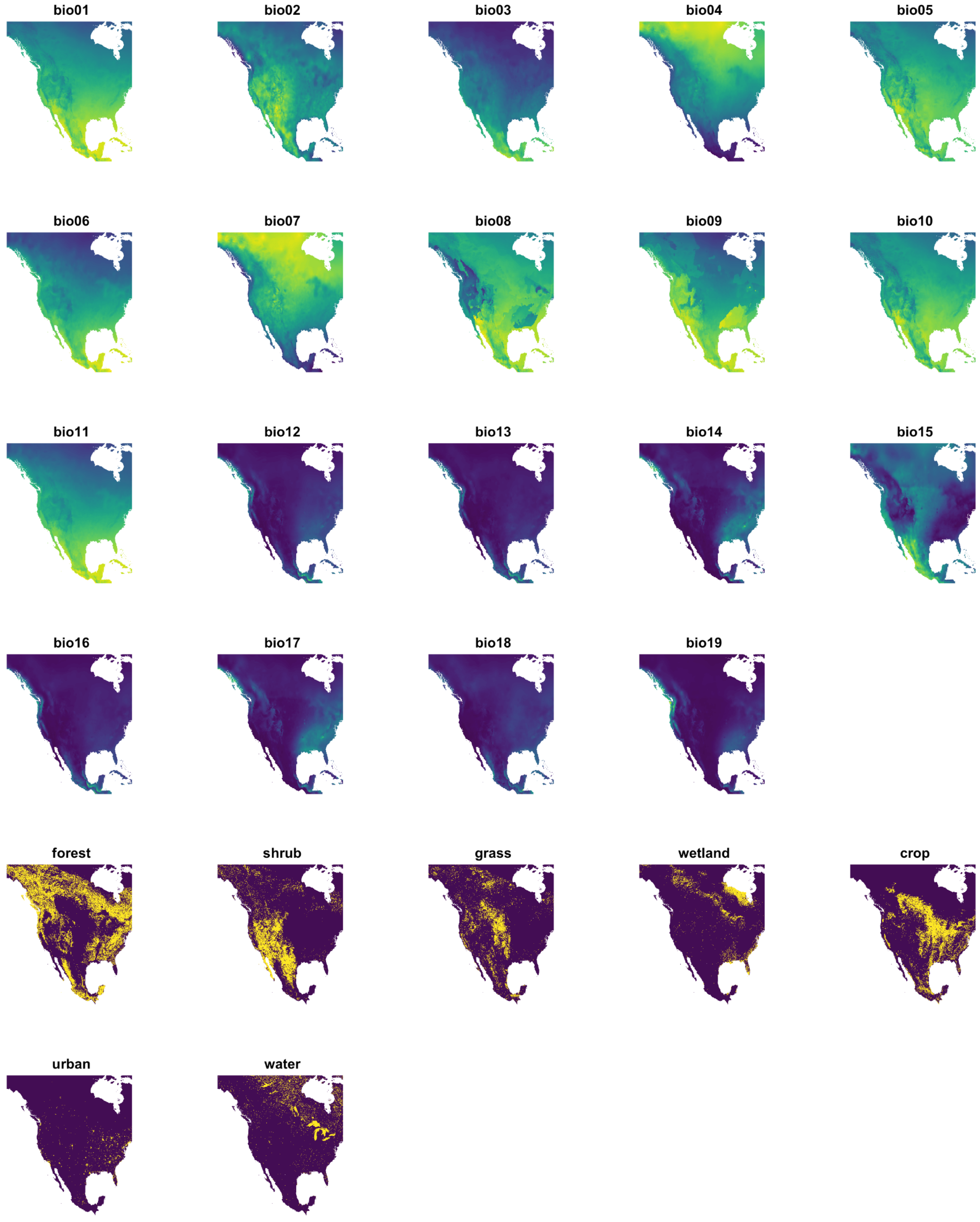


Figure S1. Maps of environmental variables used in the models, where scales are distinct across maps and reflect the range in each variable. Binary land cover variables are shown in yellow (1) and dark purple (0). See Table 1 for variable definitions.

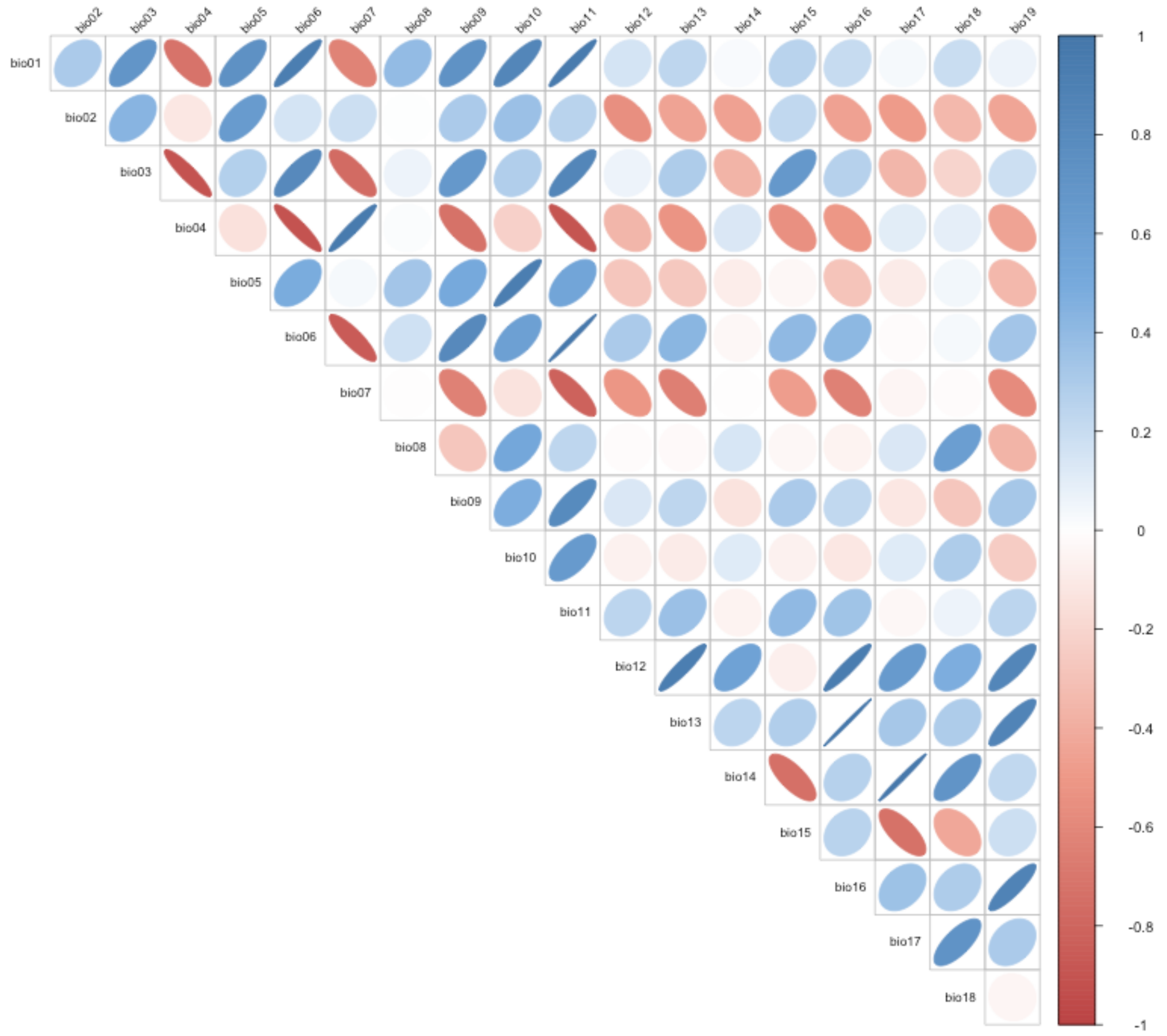


Figure S2. Correlations among bioclimatic variables for all sampling points used in the models. Correlation strength is indicated by ellipse shape (narrower indicates stronger correlation, rounder indicates weaker correlation), direction (positive points right, negative points left), and color (red is negative, blue is positive, white is zero).

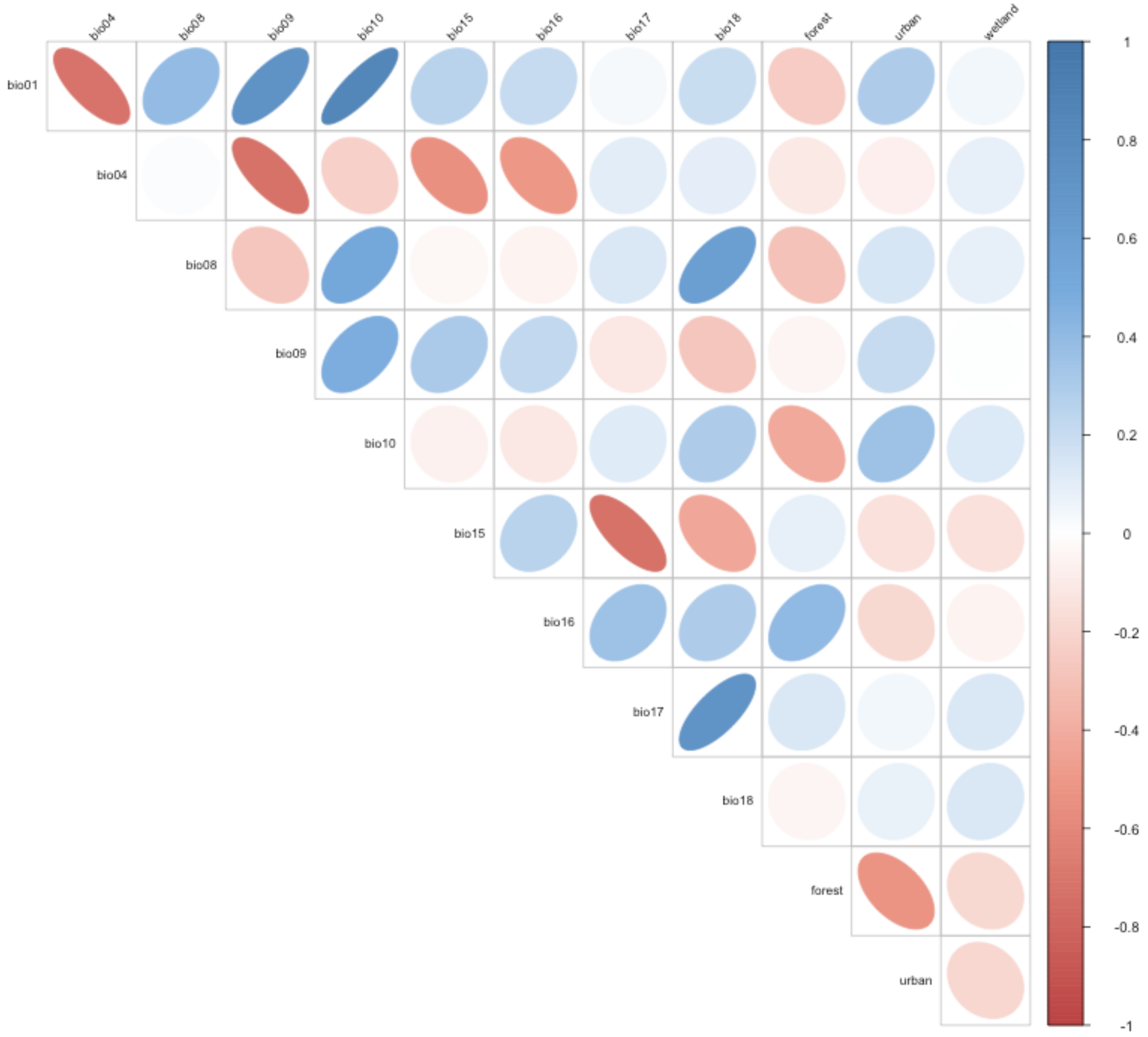


Figure S3. Correlations among a subset of bioclimatic and land cover variables for all sampling points used in the models. Correlation strength is indicated by ellipse shape (narrower indicates stronger correlation, rounder indicates weaker correlation), direction (positive points right, negative points left), and color (red is negative, blue is positive, white is zero).

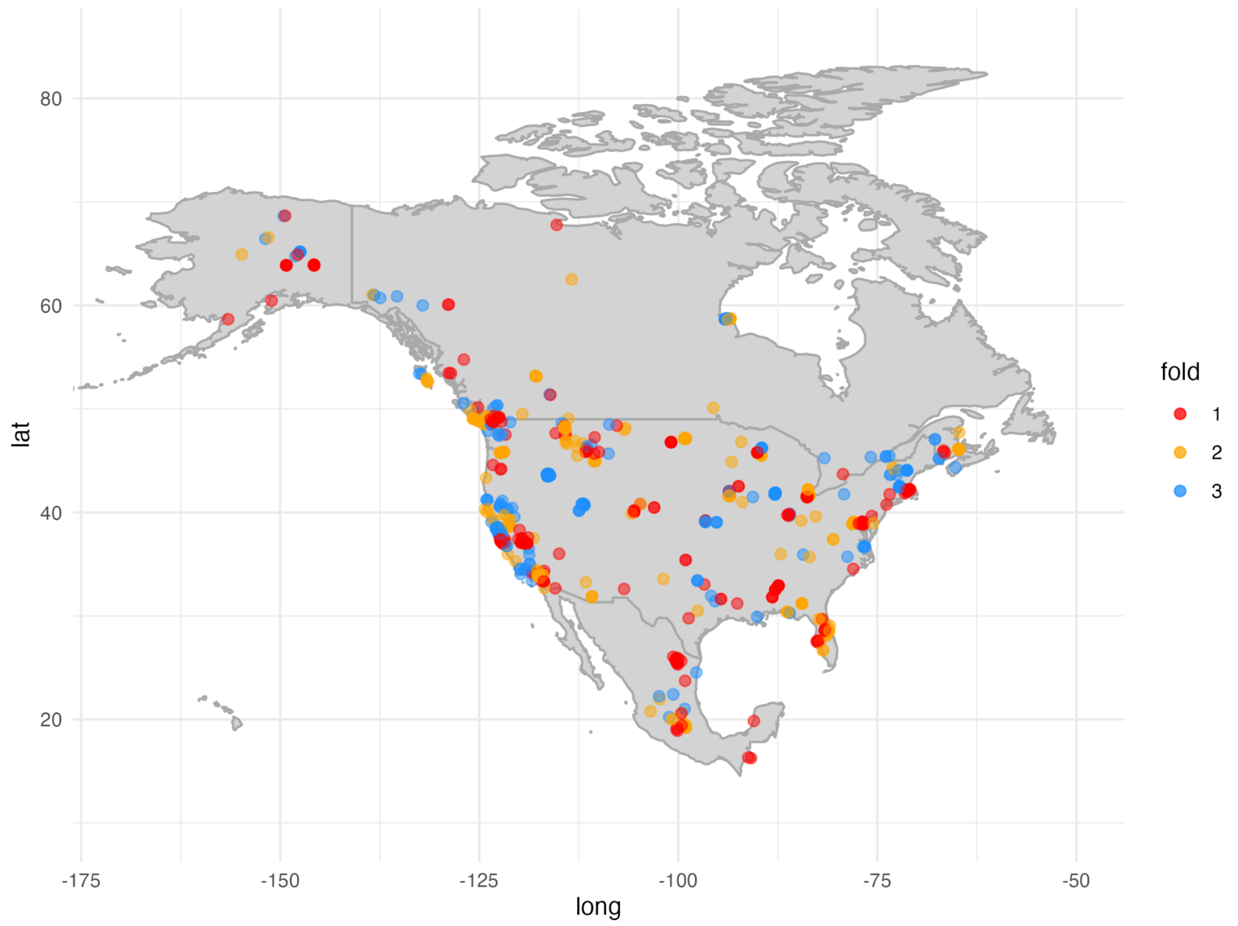


Figure S4. Spatial cross-validation fold assignments for all occurrence and background points used in the models.

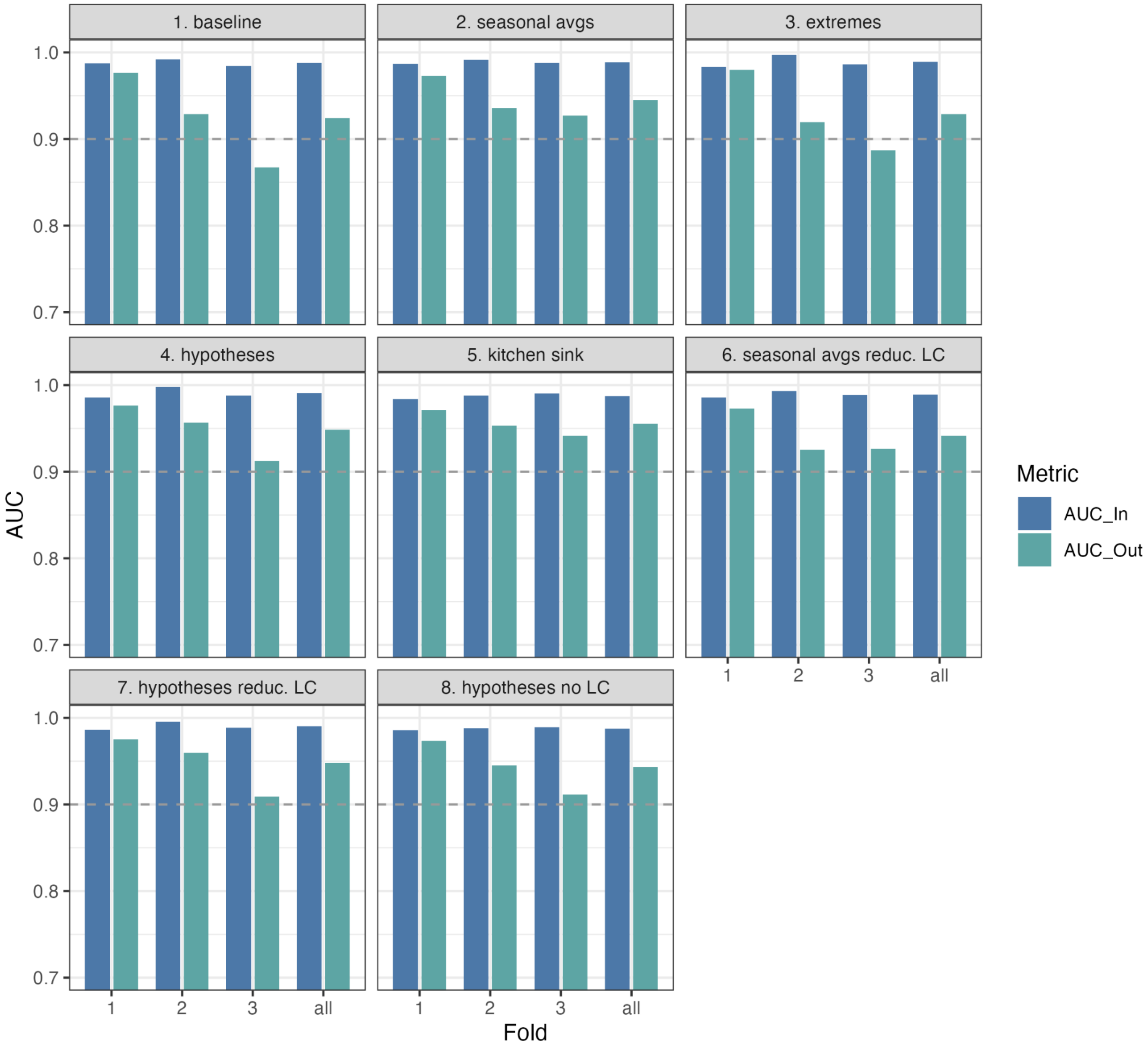


Figure S5. Model performance (area under the receiver operating characteristic curve [AUC]) for each of the 8 models, showing consistency across held-out folds (x-axis), in- and out-of-sample (blue vs. green bars), and models (panels). Note that the y-axis range is truncated to a minimum of 0.7 and a dashed line is arbitrarily shown at 0.9 for reference to distinguish performance differences among models. For model specifications, see Table 1.

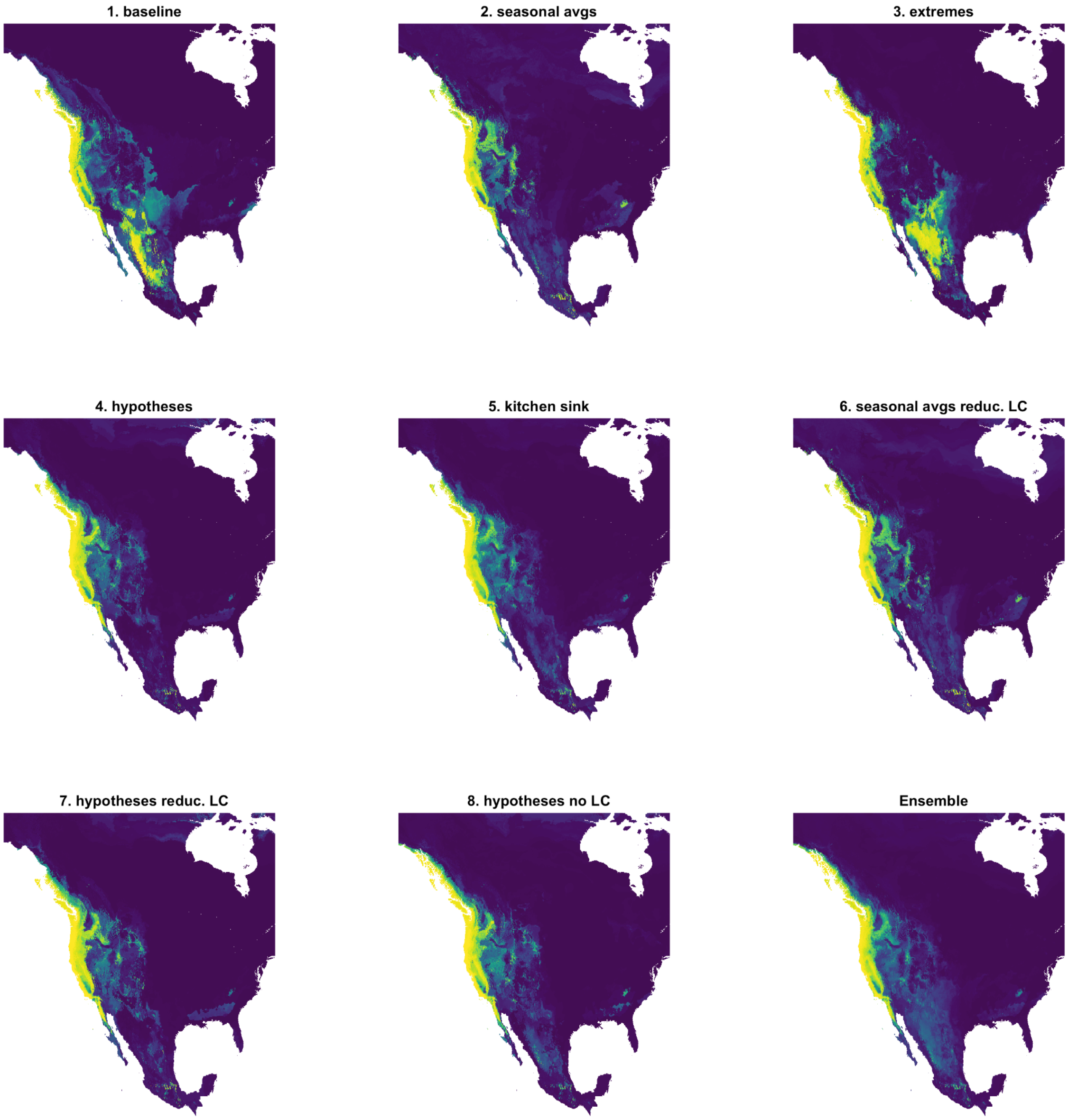


Figure S6. Model prediction maps for each individual model and the ensemble, where habitat suitability scales from zero (dark purple) to one (yellow). For model specifications, see Table 1.

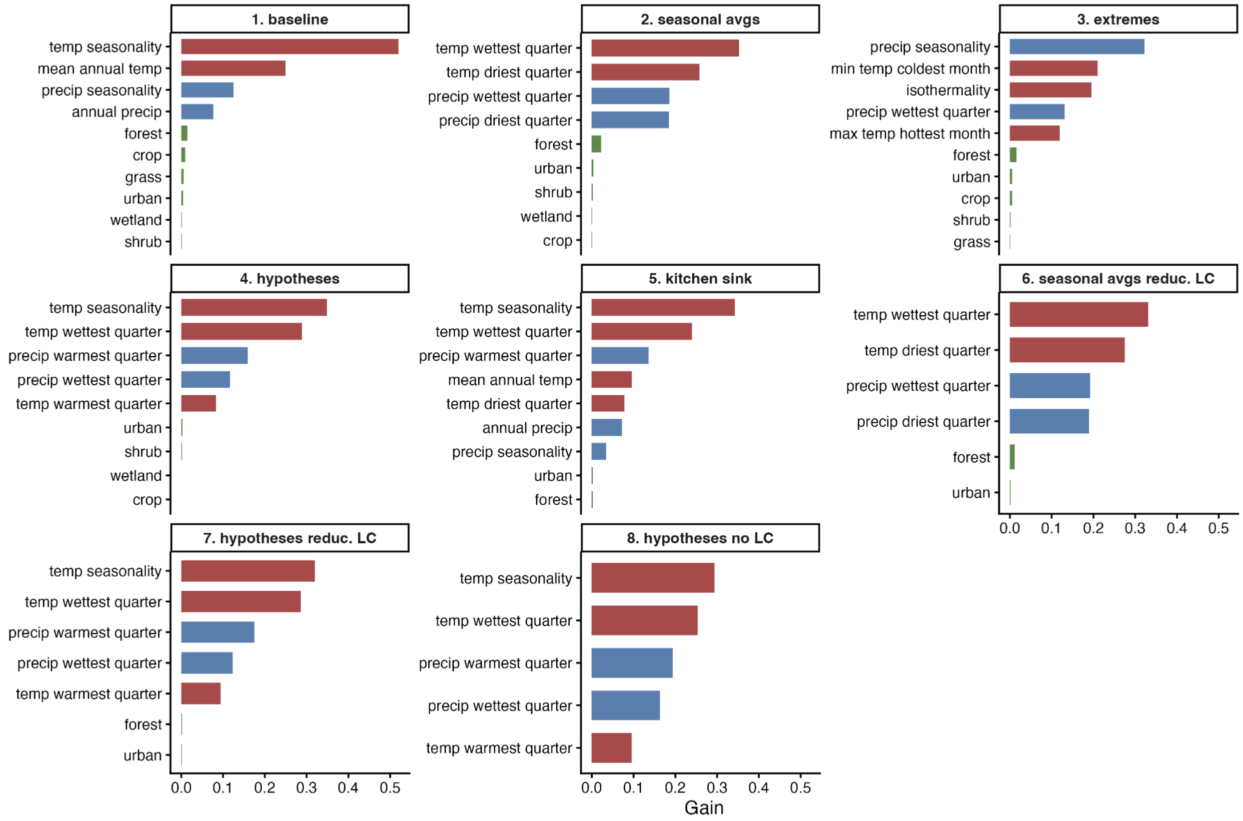


Figure S7. Variable importance based on mean gain for each model. Temperature variables are shown in red, precipitation in blue, and land cover in green. Although in some cases the rank order of bioclimatic variables differs slightly from variable importance based on SHAP (Fig. 3), bioclimatic variables were consistently more important than land cover variables when both were included in the model (models 1-7). See Table 1 for model specifications.

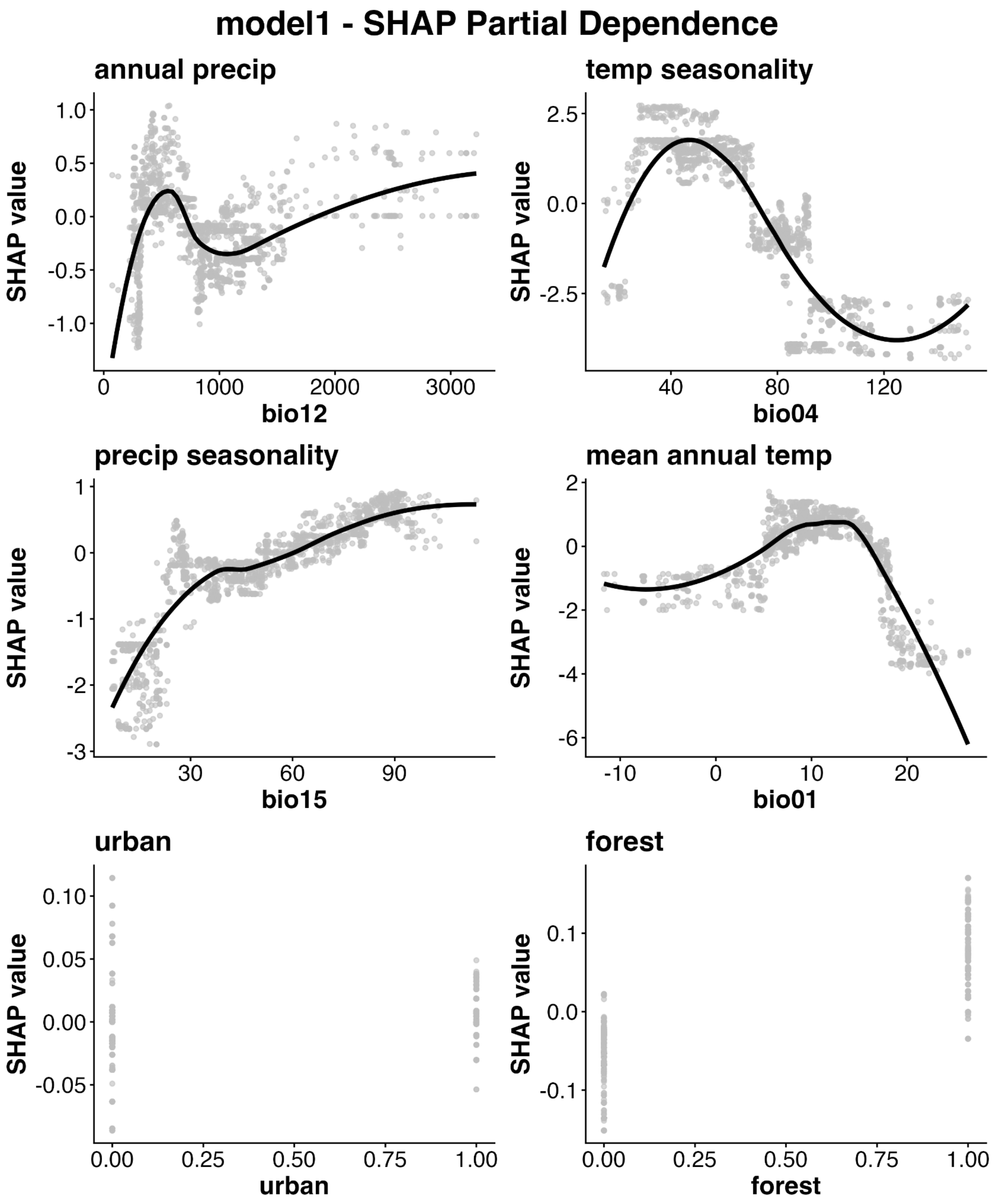


Figure S8. SHAP partial dependence plots for the top six variables in model 1.

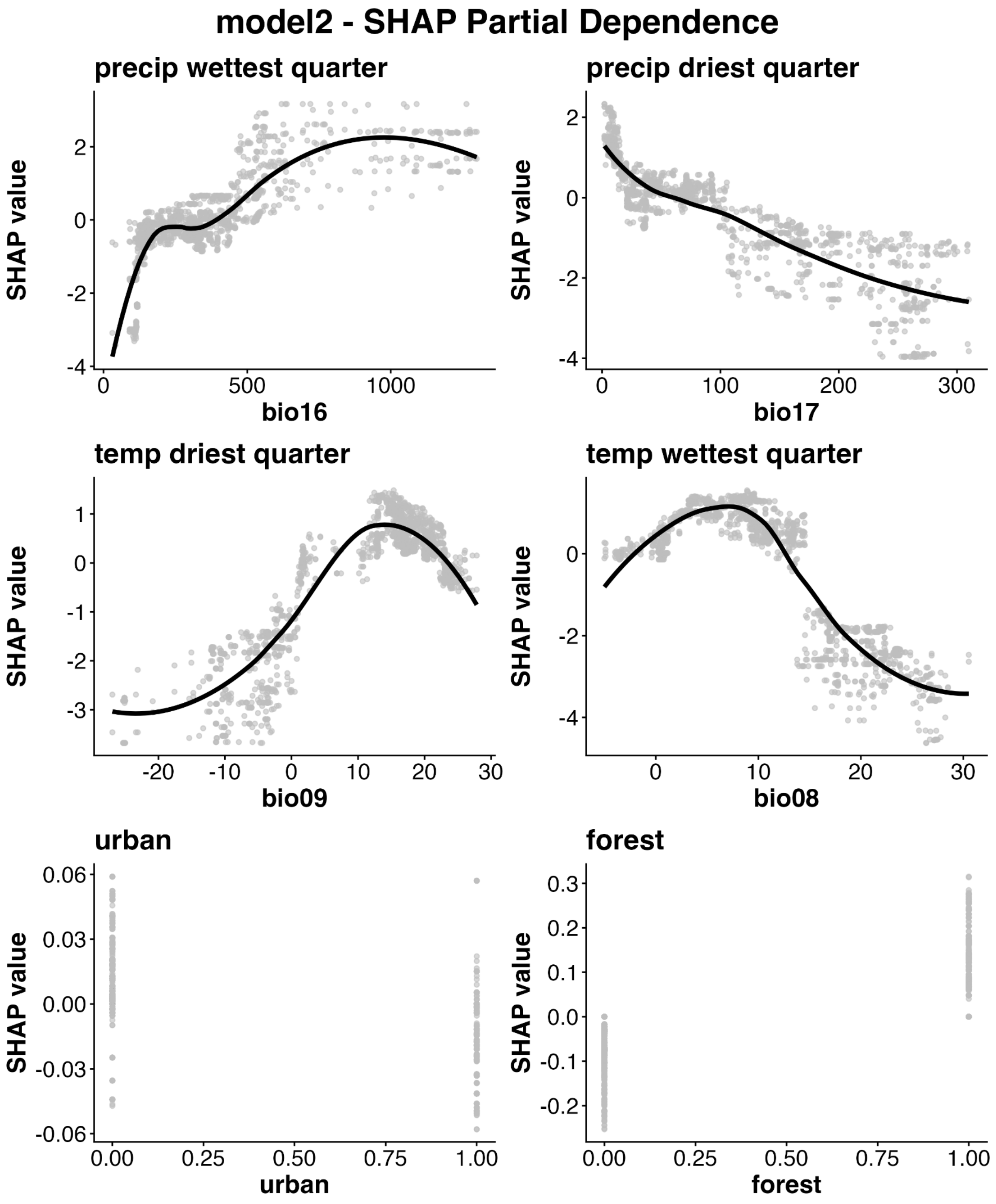


Figure S9. SHAP partial dependence plots for the top six variables in model 2.

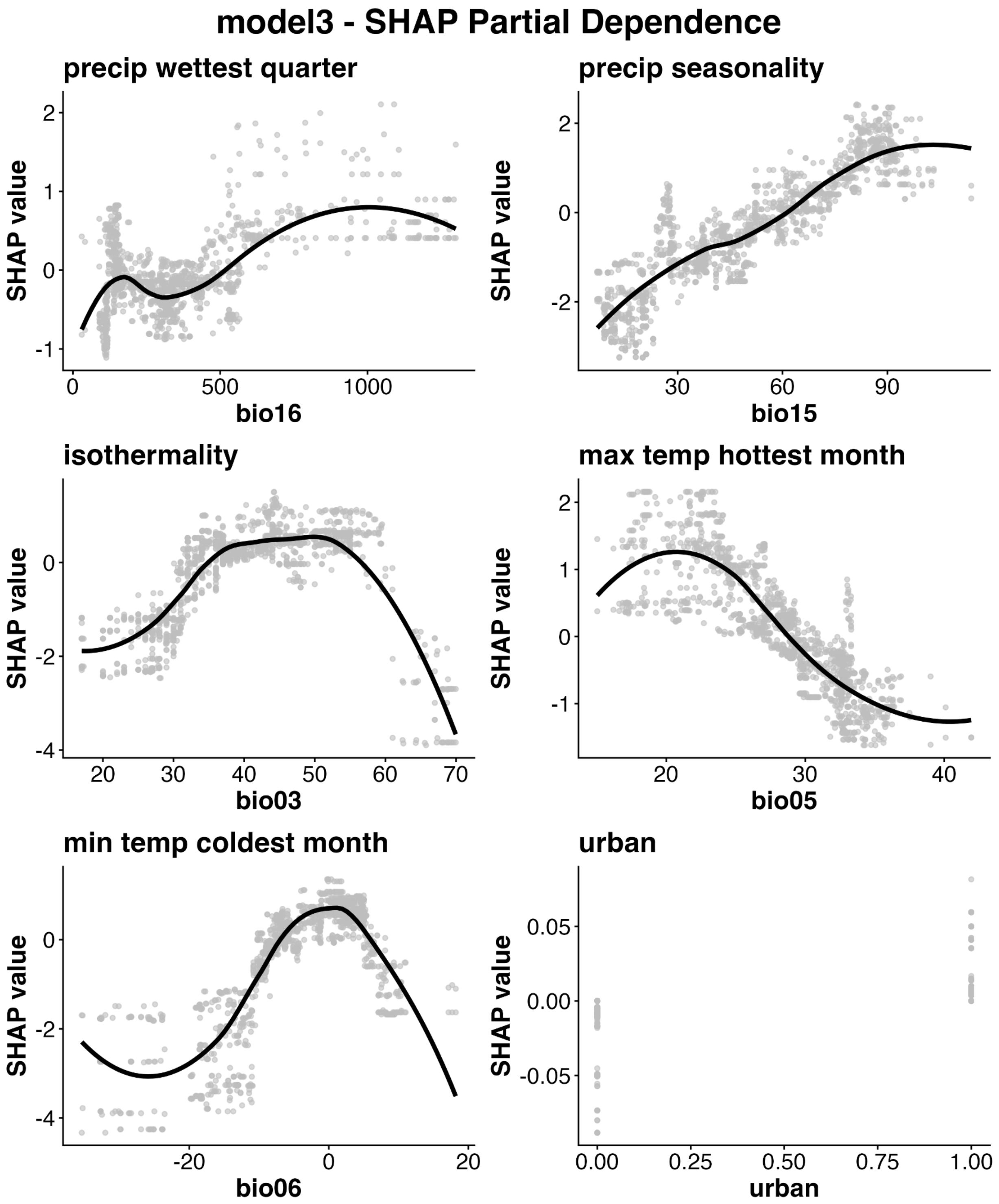


Figure S10. SHAP partial dependence plots for the top six variables in model 3.

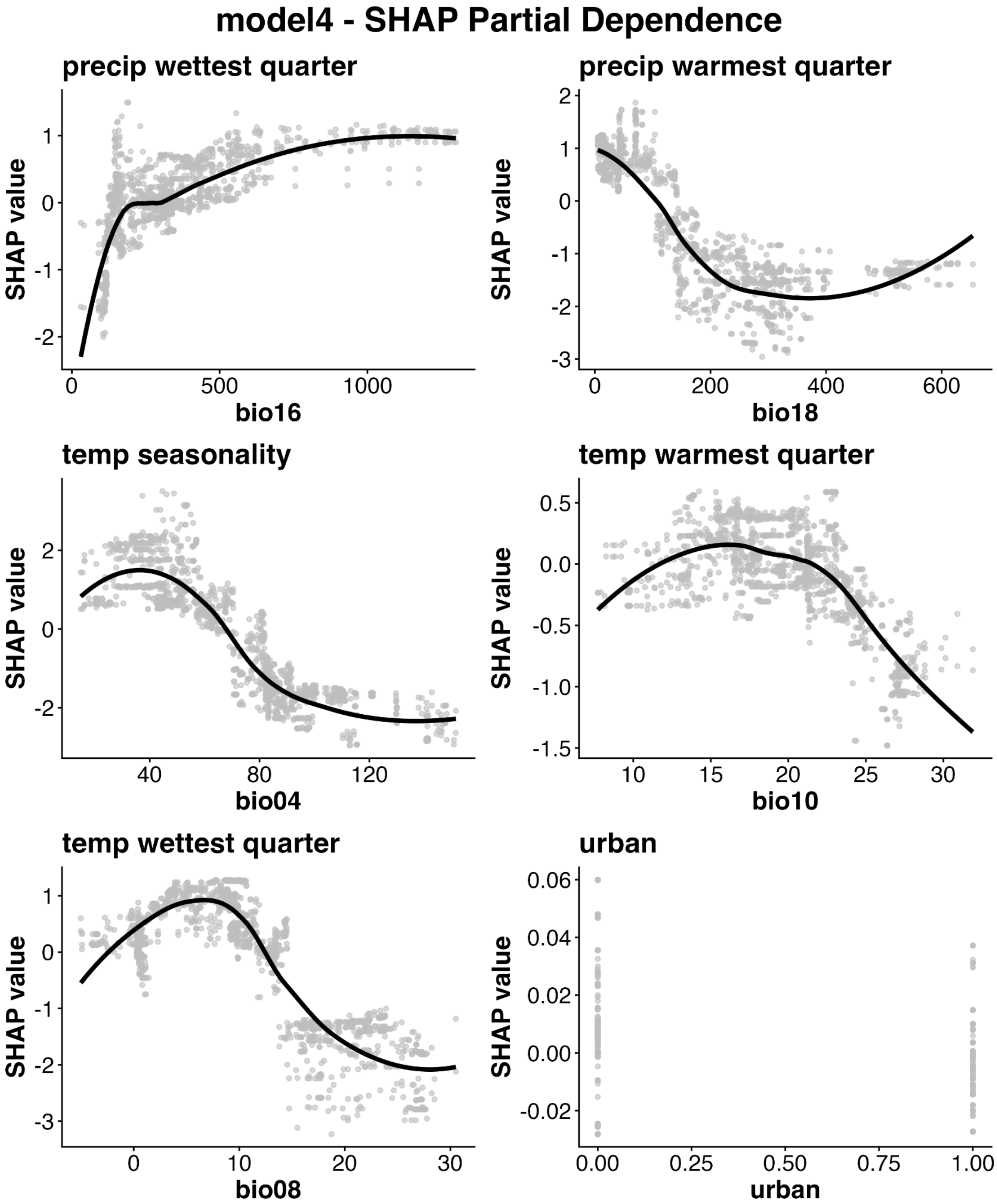


Figure S11. SHAP partial dependence plots for the top six variables in model 4.

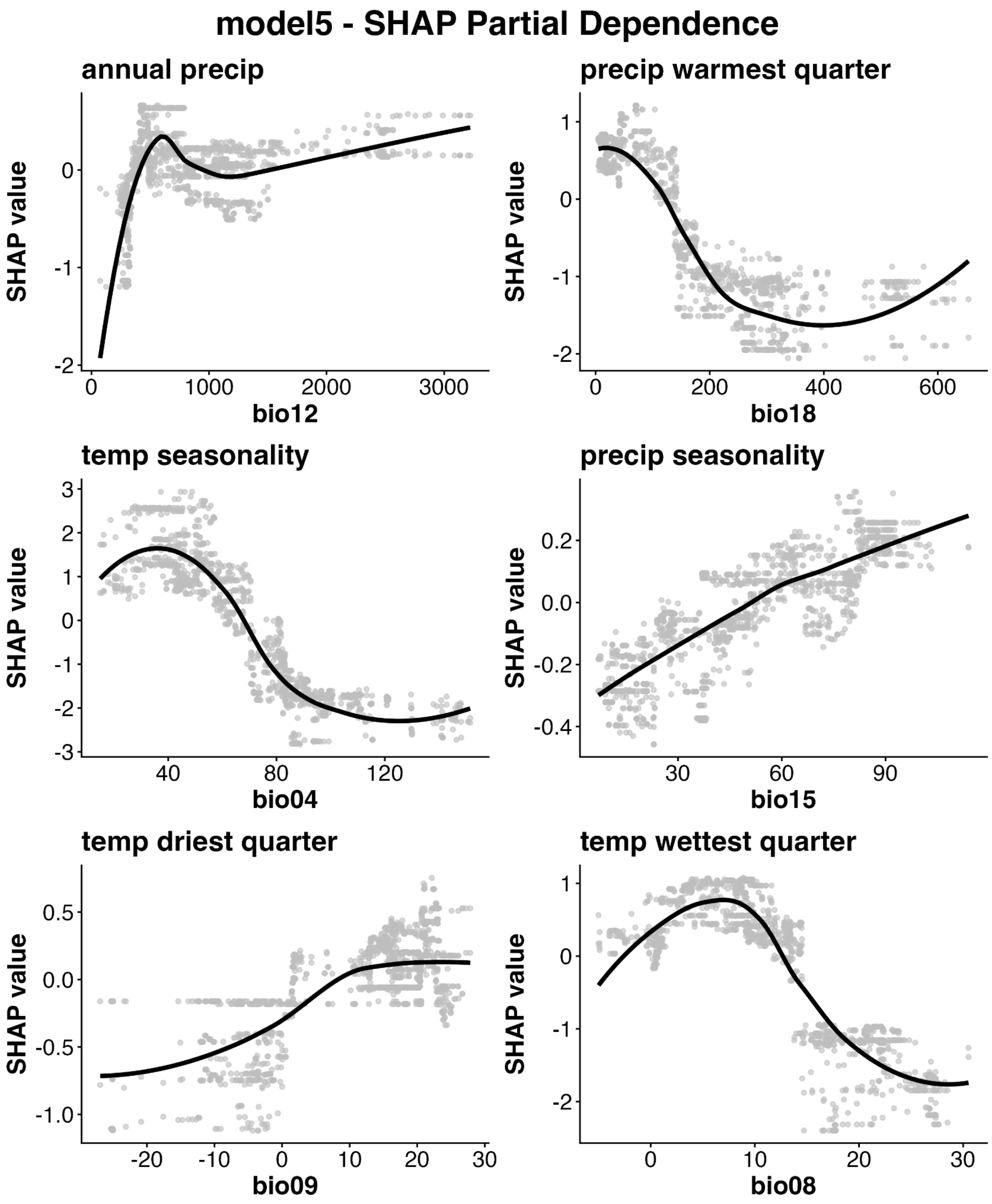


Figure S12. SHAP partial dependence plots for the top six variables in model 5.

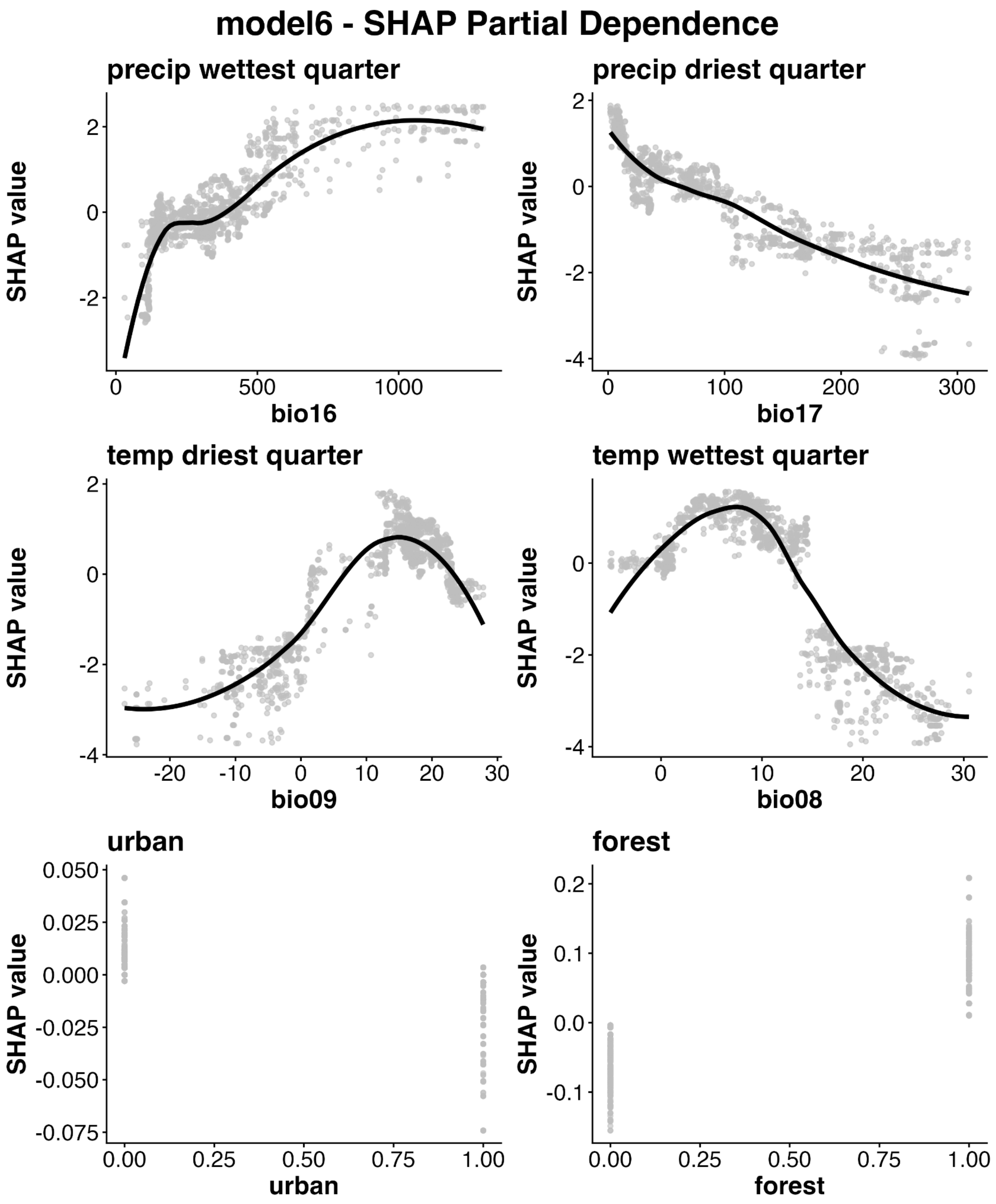


Figure S13. SHAP partial dependence plots for the top six variables in model 6.

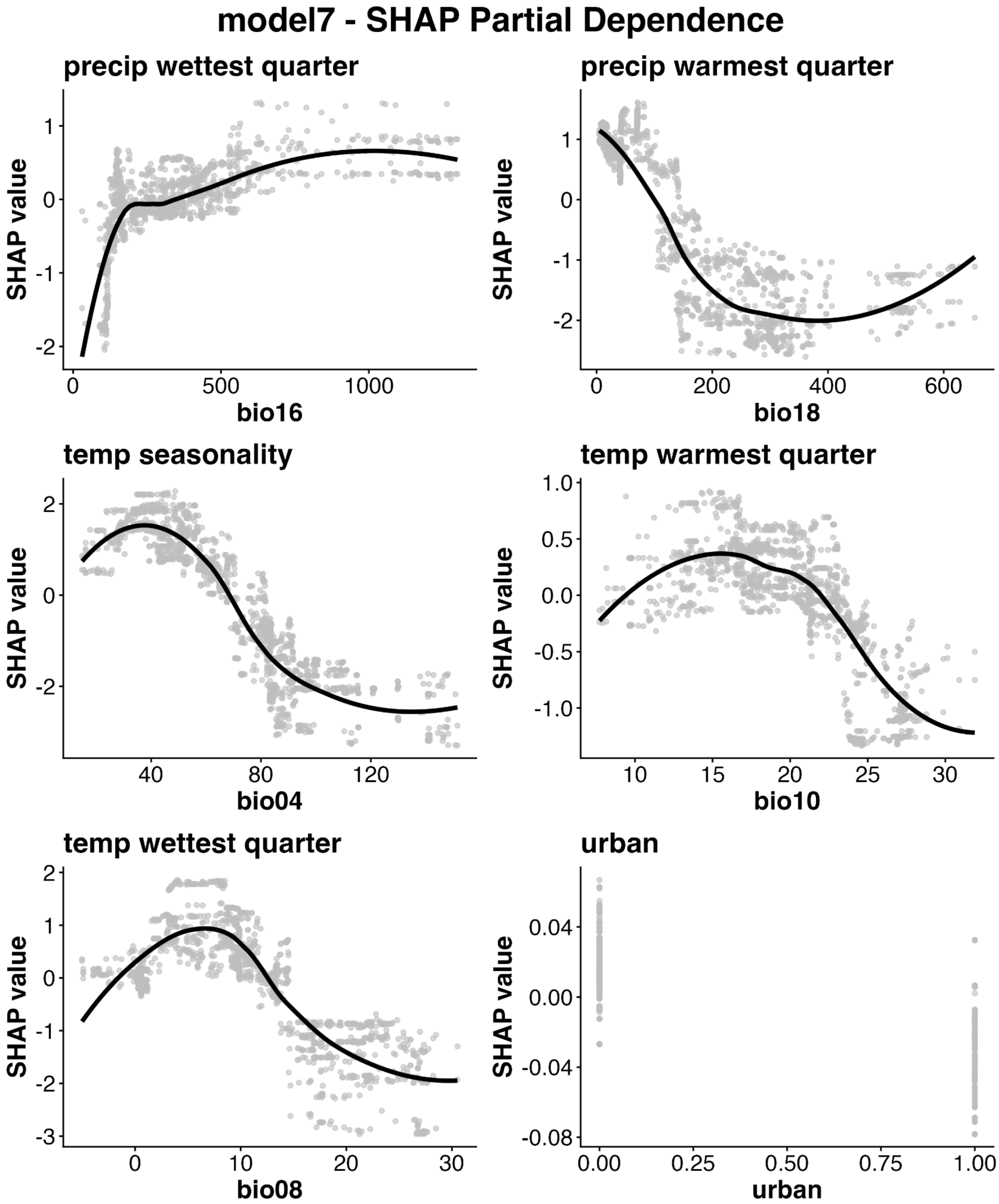


Figure S14. SHAP partial dependence plots for the top six variables in model 7.

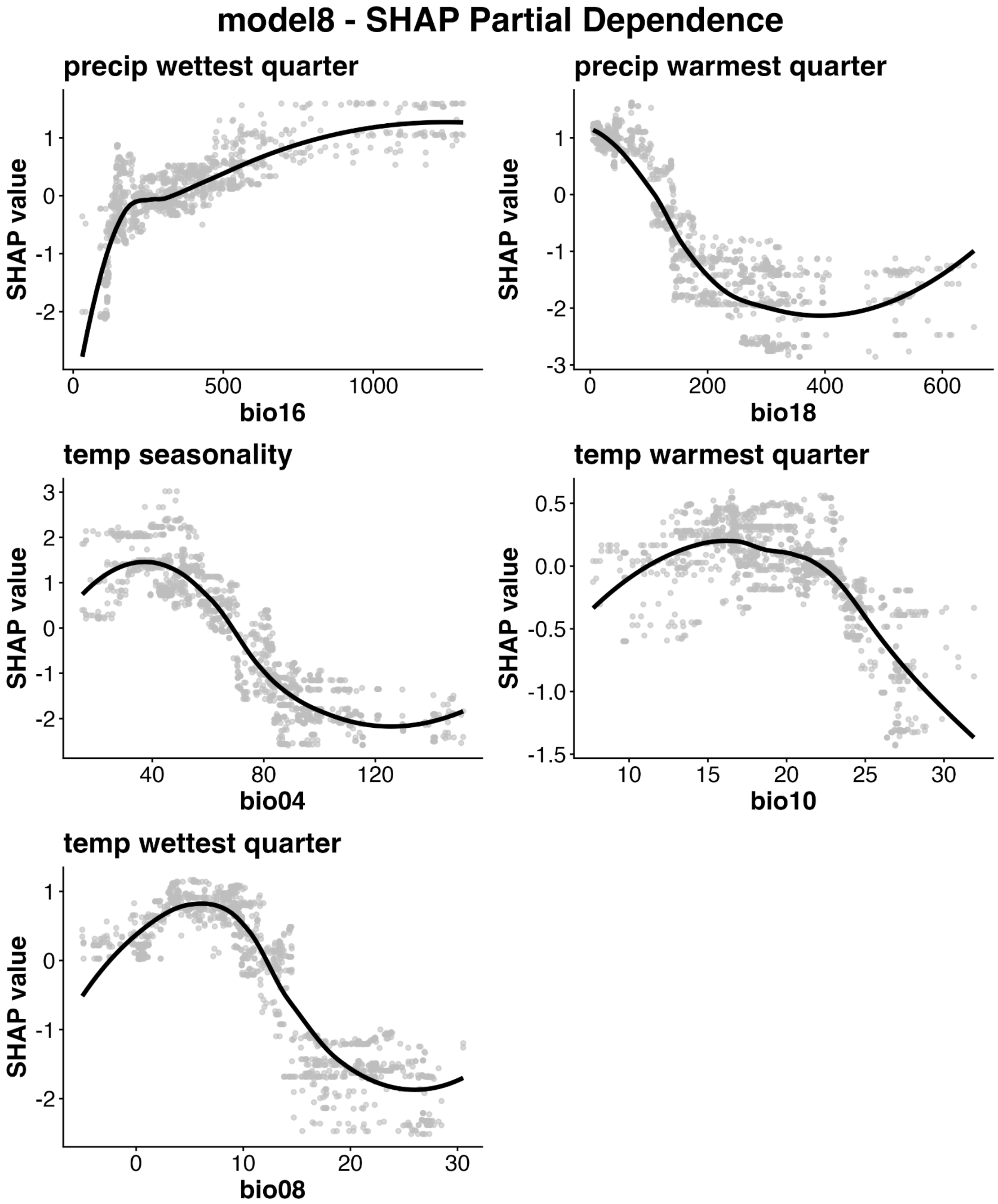


Figure S15. SHAP partial dependence plots for all variables in model 8.

#### Appendix S2. Supplementary Tables

Table S1. Occurrence and background points used to fit models, downloaded from the Global Biodiversity Information Facility (GBIF) using `rgif` package, limiting to records since 1981. Occurrence records for *Ae. sierrensis* were filtered to retain one per 1km grid cell. Background records were filtered to retain a 2:1 ratio of background to occurrence points by probabilistically subsampling on a 1km grid, for a total of 546 points. For species with fewer than 10,000 records from North America in GBIF, all records were included in the original search. For those with more than 10,000, I searched for Canada, USA, and Mexico separately to attain better geographic representation. In an effort to improve discrimination of the *Ae. sierrensis* range in the Intermountain West, USA, I specifically searched for background points in Washington, Utah, and Idaho, USA for *Ae. vexans*. I excluded points located in Hawaii (*Cx. quinquefasciatus* only) given its isolation from the rest of the species range.

| Species | Original records | Filtered records | Rationale and notes |
|---|---|---|---|
| *Aedes sierrensis* | 806 (all) | 273 | Widespread across western North America (focal species) |
| *Aedes communis* | 5624 (all) | 124 | Occurs in Alaska and northern Canada |
| *Aedes sticticus* | 9546 (all) | 134 | Occurs in eastern USA |
| *Aedes fitchii* | 1912 (all) | 22 | Occurs in Canada and USA |
| *Aedes excrucians* | 2801 (all) | 15 | Occurs in northern Canada |
| *Aedes triseriatus* | 3157 (first 3000 each from Canada, USA, Mexico) | 34 | Eastern tree hole mosquito, may have ecological similarity to *Ae. sierrensis* with distinct geographic range |
| *Aedes vexans* | 3760 (first 3000 each from Washington, Utah, and Idaho, USA) | 16 | Selected to differentiate *Ae. sierrensis* range in Intermountain West, USA |
| *Culex pipiens* | 3283 (first 3000 each from Canada, USA, Mexico) | 18 | Widespread mosquito across North America, aiming to |

| | | | |
|---|---|---|---|
| | | | sample evenly across the three countries |
| *Culex quinquefasciatus* | 5775 (first 3000 each from Canada, USA, Mexico, excluding Hawaii) | 77 | Widespread mosquito across North America, aiming to sample evenly across the three countries |
| *Culex tarsalis* | 3703 (first 3000 each from Canada, USA, Mexico) | 106 | Widespread mosquito across North America, aiming to sample evenly across the three countries |

Table S2. Like all research, this project was a learning process for me. Here, I summarize some key lessons I learned about fitting SDMs that may be useful for readers conducting related modeling approaches and applications. Many of these points are covered in other SDM literature as well (e.g., Araujo et al. 2019, Barker & MacIsaac 2022, Zurrell et al. 2020, Feng et al. 2019, Singleton et al. 2024, Lippi et al. 2023).

| |
|---|
| Model tuning: Machine learning algorithms are complex and their inner workings often opaque. Changes in settings can have important impacts on model performance and predictions. |
| - Background point selection: I found that the selection of background points could make a large difference in the final prediction maps, even though model performance, variable importance, and environment – suitability relationships were broadly similar. In particular, inclusion of background points that span beyond the focal species' occurrence points in all directions was critical for estimating reasonable environmental limits. For example, in an earlier model where I did not include background points from northern Canada, the model erroneously predicted high suitability for *Ae. sierrensis* around the Hudson Bay. |
| - Environmental variable selection: Bioclimatic variables can highly correlated with each other, but these correlations may vary geographically, so that two variables that are highly positively correlated in a region where occurrence and background points are sampled may have different correlations in other regions outside of the sampling area. As a result, inclusion of a wide range of background points (see previous point) as well as careful decisions about which environmental variables to include based on hypothesized mechanisms can have a big impact on predicted suitability outside of well-sampled areas. For example, models that included bio09 (mean temperature of the driest quarter; models 2, 5, and 6) tended to predict elevated suitability for *Ae. sierrensis* in the southeastern USA, contributing to model uncertainty in these regions (Fig. 6). |
| - Spatial cross-validation: The assignment of points to spatial folds can have important impacts on model performance and prediction outcomes, and obtaining a balance is a bit of an art. Spatial 'folds' describe spatial groupings of data points that are subsampled as a group during model training and testing—a more stringent test of model performance than simply holding out data points at random, because nearby points often share many environmental characteristics, making it easier to predict an outcome at a given point using information from nearby points. The assignment of spatial folds requires decisions about how large these spatial groupings should be and how many there should be. At the extreme, smaller spatial groupings begin to approximate random subsampling (inflating model performance but potentially losing generality through overfitting), whereas larger groupings can risk omitting key portions of environmental space (reducing out-of-sample model performance and potentially also losing generality). |
| - Order of operations: Bayesian hyperparameter optimization (BO), spatial cross- |

| | |
|---|---|
| | validation (CV), and final model fitting and interpretation can occur in many possible orderings, which can have impacts on the results. Ideally, BO occurs within each spatial CV loop so that parameters are optimized for that particular dataset, then averaged for the final model fit in which data are held out in a non-spatially structured way to train the final model. |
| - | As Lippi et al. (2023) note: “Variable importance is influenced by nearly every step of the SDM building process, such as choice of data products, scale of analysis, collinearity reduction techniques, and choice of SDM algorithm.” Cianci et al. (2015) further support this conclusion for three mosquito species native to the Netherlands, where different model algorithms produced similar prediction maps and accuracy but differing rank order of variable importance. Singleton et al. (2024) similarly find that for *Biomphalaria* snails that transmit schistosome parasites, different model algorithms produced models with similar accuracy but distinct suitability maps. |

### Appendix S3. ODMAP Protocol

Temperature and precipitation shape the distribution of *Aedes sierrensis* in western North America

Erin Mordecai

2026-06-18

---

#### Overview

***Authorship***

Contact :

Study link: To be added after publication

***Model objective***

Model objective: Inference and explanation, mapping and interpolation

***Focal Taxon***

Focal Taxon: *Aedes sierrensis* (Ludlow, 1905)

***Location***

Location: North America (USA, Canada, Mexico)

***Scale of Analysis***

Spatial extent: longitude: -168.00, -52.00, latitude: 14.5, 83.1

Spatial resolution: 1 km

Temporal extent: 1981-2026

Temporal resolution: NA

Boundary: political

***Biodiversity data***

Observation type: citizen science (GBIF)

Response data type: presence-only

***Predictors***

Predictor types: climatic, habitat

***Hypotheses***

Hypotheses: *Aedes sierrensis* distribution is limited to highly seasonal environments with cool wet seasons and hot dry seasons (e.g., Mediterranean climates) and vegetated areas with trees that can serve as immature habitat, and further limited by extreme summer highs and winter lows and sufficient rainfall to sustain several months per year in which the species' preferred water-filled tree hole immature habitat is available.

***Assumptions***

Model assumptions: Stable range (quasi-equilibrium), representative sampling of focal species, background points (occurrence records of other mosquito species) accurately represent spatial sampling biases, no uncertainty or error in predictor or response data, correlative relationships plausibly related to underlying mechanisms

***Algorithms***

Modelling techniques: brt (XGBoost)

Model complexity: Model algorithm allows for flexible, nonlinear, interactive relationships among predictor variables; uses Bayesian hyperparameter optimization (BO) to tune model fit; uses spatial blocked cross-validation to reduce overfitting and spatial leakage; uses statistics such as AUC to evaluate model performance and SHAP to evaluate variable contributions and environmental response functions.

Model averaging: Model averaging across candidate models with different environmental covariates was used to create an ensemble and to evaluate locations of model disagreement, as high-uncertainty portions of the range

***Workflow***

Model workflow: A set of candidate XGBoost model specifications (each containing different subsets of environmental predictors) were fit to predict the probability of focal species occurrence against background points (interpreted as habitat suitability). Model fits started with Bayesian hyperparameter optimization and model fitting across separate held out spatially stratified folds, to evaluate model performance and consistency across folds. A final model was fit was performed using all data to generate the final prediction map. Model evaluation was performed using area under the receiver operating characteristic curve (AUC) and log loss across folds and in and out of sample, as well as Shapley statistics for evaluating predictor feature importance and environmental responses (including visual scrutiny of both single- and two-variable partial dependence plots). Final model prediction maps were checked against input response data to ensure agreement and spatial coverage.

***Software***

Software: R version 4.5.2 (2025-10-31); packages: caret, corrplot, data.table, dismo, dplyr, forcats, future.apply, ggnewscale, ggplot2, mltools, mgcv, patchwork, pdp, PerformanceAnalytics, pROC, purrr, raster, rBayesianOptimization, RColorBrewer, rgbif, rnaturalearth, rsample, sf, SHAPforxgboost, spatialsample, terra, tidyr, tidytext, tidyverse, vip, and xgboost.

Environmental data extraction was performed in Google Earth Engine using the Google Colab python interface.

Code availability: To be deposited upon submission

Data availability: To be deposited upon submission

### Data

***Biodiversity data***

Taxon names: Focal species: *Aedes sierrensis*; Other species used as background points: *Aedes communis, Aedes sticticus, Aedes fitchii, Aedes excrucians, Aedes triseriatus, Culex tarsalis, Culex pipiens, Culex quinquefasciatus*

Taxonomic reference system: As provided in GBIF records

Ecological level: species

Data sources: GBIF.org, accessed on June 23, 2026

Sampling design: Convenience sampling (citizen science)

Basis of observation: human observation, preserved specimen, or material sample

Sample size: N = 273 *Aedes sierrensis* occurrence points after spatial filtering

Absence data: N = 546 background points (occurrence records for other mosquito species) after spatial filtering and probabilistic subsampling

Background data: Background points were sampled from the listed species to cover the broad geographic extent of USA, Canada, and Mexico, excluding Hawaii, as described in Table S1

***Predictor variables***

Predictor variables: Bioclimatic variables from WorldClim BioClim v1; land cover data from NLCD National Land Cover of North America 2020

Data sources: All data accessed on June 24, 2026

Spatial extent: longitude: -168, -52, latitude: 14.5, 83.1

Spatial resolution: Bioclim: 1km; NLCD: 30m

Coordinate reference system: Bioclim: WGS 84 geographic coordinate system, EPSG Code: 4326; NLCD: EPSG Code: 5070

Temporal extent: Bioclim: 1950-2000; NLCD: 2019-2021

***Transfer data***

Spatial extent: NA, NA, NA, NA (xmin, xmax, ymin, ymax)

Spatial resolution: NA

Temporal extent: NA

### Model

***Multicollinearity***

Multicollinearity: Examined correlations among predictors and developed a set of candidate models, each containing a subset of predictor variables that: (a) capture variation in hypothesized temperature, precipitation, and land cover drivers, and (b) have correlations lower than 0.7 absolute value.

***Model settings***

See Methods > Model fitting > XGBoost modeling methods for final model specifications and settings.

***Model estimates***

Coefficients: NA

***Analysis and Correction of non-independence***

Spatial autocorrelation: Spatial block cross-validation

### Assessment

***Performance statistics***

Performance on training data and out-of-sample testing data: AUC, logloss

***Plausibility check***

Response shapes: Variable importance and partial dependence plots using SHAP scores; Prediction map: concurrence with recorded occurrence and background points and plausible geographic distribution
